# A Missense Point Mutation in Nerve Growth Factor (NGF^R100W^) Results in Selective Peripheral Sensory Neuropathy

**DOI:** 10.1101/784660

**Authors:** Wanlin Yang, Kijung Sung, Wei Xu, Maria J Rodriguez, Andrew C. Wu, Sarai A. Santos, Savannah Fang, Rebecca K. Uber, Stephanie X. Dong, Brandon C. Guillory, Xavier Orain, Jordan Raus, Corrine Jolivalt, Nigel Calcutt, Robert A. Rissman, Jianqing Ding, Chengbiao Wu

## Abstract

A missense point mutation in nerve growth factor (NGF^R100W^) is associated with hereditary sensory autonomic neuropathy V (HSAN V), originally discovered in a Swedish family. These patients develop severe loss of perception to deep pain but with apparently normal cognitive functions. To better understand the disease mechanism, we have generated the first NGF^R100W^ knockin mouse model of HSAN V. Mice homozygous for the NGF^R100W^ mutation (NGF^fln/fln^) showed significant structural deficits in intra-epidermal nerve fibers (IENFs) at birth. These mice had a total loss of pain perception at ∼2 months of age and they often failed to survive to full adulthood. Heterozygous mice (NGF^+/fln^) developed a progressive degeneration of small sensory fibers both behaviorally and functionally: they showed a progressive loss of IENFs starting at the age of 9 months accompanied with progressive loss of perception to painful stimuli such as noxious temperature. Quantitative analysis of lumbar 4/5 dorsal root ganglia (DRG) revealed a significant reduction in small size neurons positive for calcitonin gene-related peptide, while analysis of sciatic nerve fibers revealed the mutant NGF^+/fln^ mice had no reduction in myelinated nerve fibers. Significantly, the amount of NGF secreted from fibroblasts were reduced in heterozygous and homozygous mice compared to their wild-type littermates. Interestingly, NGF^+/fln^ showed no apparent structural alteration in the brain: neither the anterior cingulate cortex nor the medial septum including NGF-dependent basal forebrain cholinergic neurons. Accordingly, these animals did not develop appreciable deficits in tests for central nervous system function. Our study provides novel insights into the selective impact of NGF^R100W^ mutation on the development and function of the peripheral sensory system.

## INTRODUCTION

Nerve growth factor (NGF) is a member of the neurotrophin family that includes brain-derived neurotrophic factor (BDNF), neurotrophin 3 (NT-3) and neurotrophin 4 (NT-4) (Chao, 2003; Chao and Hempstead, 1995; Huang and Reichardt, 2001; Levi-Montalcini, 1987, 2004; Levi-Montalcini et al., 1995). These trophic factors act through two distinct receptors, Trk, the 140 kD tyrosine receptor kinase (TrkA for NGF; TrkB for BDNF, NT-3; TrkC for NT-4) and the 75 kD neurotrophin receptor (p75^NTR^) to transmit signals in responsive neurons (Bothwell, 1995; Chao and Hempstead, 1995; Kaplan and Miller, 1997). The trophic function of neurotrophins is largely mediated by Trk, while p75^NTR^ has a more diverse effects on survival, differentiation and death of neurons (Casaccia-Bonnefil et al., 1998; Chao and Hempstead, 1995; Nykjaer et al., 2005).

NGF exerts potent trophic actions on sensory and sympathetic neurons of the peripheral nervous system (PNS)(Hamburger and Levi-Montalcini, 1949) and also regulates the trophic status of striatal and basal forebrain cholinergic neurons (BFCNs) of the central nervous system (CNS)(Conover and Yancopoulos, 1997; Kew et al., 1996; Lehmann et al., 1999; Levi-Montalcini and Hamburger, 1951; Li and Jope, 1995; Svendsen et al., 1994). Given its robust trophic effects, NGF has been investigated for therapeutic properties for treating both PNS and CNS diseases. For example, NGF was explored for treating/preventing degeneration of BFCNs in Alzheimer’s disease (AD) (Blesch and Tuszynski, 1995; Cuello et al., 2010; Eriksdotter Jonhagen et al., 1998; Hefti, 1994; Knusel and Gao, 1996; Koliatsos, 1996; Mufson et al., 2008; Olson, 1993; Rafii et al., 2014; Schindowski et al., 2008; Schulte-Herbruggen et al., 2008; Scott and Crutcher, 1994; Williams et al., 2006). However, some significant pain issues associated with administration of recombinant NGF such as back pain, injection site hyperalgesia, myalgia, etc led to the termination of these trials(Eriksdotter Jonhagen et al., 1998; Hefti, 1994; Knusel and Gao, 1996; Koliatsos, 1996; Olson, 1993; Scott and Crutcher, 1994). Other clinical efforts using NGF for treating diabetic neuropathies, HIV-induced peripheral neuropathies were also terminated due to the extreme side effects of pain(Apfel, 2002; Apfel et al., 1998; Hellweg and Hartung, 1990; Lein, 1995; McArthur et al., 2000; Pradat, 2003; Quasthoff and Hartung, 2001; Rask, 1999; Schifitto et al., 2001; Unger et al., 1998; Walwyn et al., 2006). These adverse effects including severe pain associated with NGF administration has proven to be a significant roadblock in advancing NGF to therapies.

Indeed, NGF has also been recognized as a potent mediator of pain (Chuang et al., 2001; Lewin and Mendell, 1993; Lewin et al., 1993; Watanabe et al., 2008). This is further supported by genetic and clinical evidence demonstrating that both TrkA and p75^NTR^-mediated signaling contributes to NGF-induced hyper-sensitization. For example, hereditary sensory and autonomic neuropathy type IV (HSAN IV) is resulted from recessive mutations in TrkA, these patients display pain insensitivity as well as mental retardation (Indo, 2001, 2002). Furthermore, many TrkA downstream effectors have also been implicated in NGF-mediated nociception: Pharmacological Inhibition of either the extracellular signal-regulated kinases (Erk) or phosphoinositide 3-kinase (PI3K) attenuates NGF-induced hyperalgesia (Aley et al., 2001; Dai et al., 2002; Zhuang et al., 2004). Activation of Phospholipase C (PLCγ) by NGF also potentiates nociceptive ion channels leading to hyperalgesia (Chuang et al., 2001). p75^NTR^ is also involved in NGF-induced hyperalgesia. For example, injecting a p75^NTR^ neutralizing antibody blocked NGF-induced hyperalgesia and NGF-mediated sensitization of action potentials in sensory neurons (Watanabe et al., 2008; Zhang and Nicol, 2004). Ceramide, a p75 downstream effector, is known to increase the number of action potentials in sensory neurons (Zhang et al., 2002, 2006). Nerve injury, axotomy, seizure, or ischemia can all cause an increase in both the expression and axonal transport of p75^NTR^, thereby contributing to nociception (Zhou et al., 1996) (Roux et al., 1999). p75^NTR^ downstream signaling cascades were responsible for mechanical hyperalgesia following NGF injection (Khodorova et al., 2013). Therefore, both TrkA and p75^NTR^ receptor(s)-mediated signaling pathways play an important role in the pain signaling and function of NGF, although their relative contribution is yet to be defined.

The discovery of NGF mutations in human patients further highlights the importance of NGF in nociception. A homozygous mutation in the NGF gene [680C>A]+[681_682delGG] has been linked to hereditary sensory neuropathy in a consanguineous Arab family(Carvalho et al., 2011). The affected individuals were completely unable to perceive pain, did not sweat, could not discriminate temperature(Carvalho et al., 2011). In addition, these patients had a chronic immunodeficiency, and they were intellectually disabled (Carvalho et al., 2011).

In a second case, a missense mutation in NGF (661C>T) was discovered in patients in consanguineous Swedish families who suffered from severe loss of deep pain, bone fractures and joint destruction(Carvalho et al., 2011; Einarsdottir et al., 2004). The disorder was classified as HSAN V (Online Mendelian Inheritance in Man (OMIM) # 608654). This particular mutation leads to a substitution of tryptophan (W) for arginine (R) at position 211 in the proNGF polypeptide (pro-NGF^R221W^) and at position 100 in the mature protein (NGF^R100W^) (Einarsdottir et al., 2004). Interestingly, unlike HSAN IV patients with recessive TrkA mutation(s) that develop pain insensitivity as well as mental retardation, HSAN V patients suffer from selective loss of pain sensation with normal cognitive function, suggesting that the NGF^R100W^ mutation causes selective loss of pain function, but likely with intact trophic function (Einarsdottir et al., 2004; Minde et al., 2009; Minde et al., 2004b; Minde, 2006; Perini et al., 2016; Sagafos et al., 2016). Therefore, this NGF^R100W^ mutant can provide an important tool to uncouple the trophic function of NGF from its nociceptive actions(Capsoni, 2014; Capsoni et al., 2011; Cattaneo and Capsoni, 2019; Sung et al., 2018; Sung et al., 2019; Testa et al., 2019; Testa et al., 2018; Yang et al., 2018).

Previous studies have revealed that mutation(s) at the R100 position might disrupt the processing of proNGF to mature NGF (Larsson et al., 2009). We also previously examined the binding and signaling properties of the mature form of naturally occurring mutant NGF (NGF^R100W^) and discovered that NGF^R100W^ retains binding and signaling through TrkA to induce trophic effects but failed to bind or activate p75^NTR^ (Sung et al., 2018). Consistent with our studies, other mutation(s) at the R100 position (NGF^R100E^ and NGF^P61SR100E^) didn’t affect NGF binding to TrkA, while it abolished NGF binding to p75^NTR^ (Capsoni et al., 2011; Covaceuszach et al., 2010). Together, these provide strong support a role of p75^NTR^ in NGF-induced pain function and failure to activate p75^NTR^ signaling is likely the cause for loss of pain in HSAN V patients(Capsoni, 2014; Capsoni et al., 2011; Cattaneo and Capsoni, 2019; Sung et al., 2018; Sung et al., 2019; Testa et al., 2019; Testa et al., 2018; Yang et al., 2018).

HSAN V is an extremely rare disease that makes it almost impossible to perform systematic study of NGF^R100W^ in terms of its mechanism and function *in vivo*. Given that NGF is highly conserved between mice and human, mouse models of HSAN V carrying NGF^R100W^ provide an important vehicle to carefully examine the effects of NGF^R100W^ on development/ maintenance of CNS and PNS *in vivo* (Testa et al., 2019; Testa et al., 2018; Yang et al., 2018). Using the knock-in mouse NGF^R100W^ model of HSAN V, we previously reported that mice carrying homozygous NGF^R100W^ alleles (NGF^fln/fln^) displayed normal embryonic development of major organs (heart, lung, liver, kidney, and spleen) showed normal gene expression of either TrkA or p75^NTR^ receptor(Yang et al., 2018). Interestingly, they exhibited extremely low counts of IENFs at birth(Yang et al., 2018).

In the present study, we characterized the impact of NGF^R100W^ on both the brain and peripheral sensory nerves, both structurally and functionally, using our knock-in mouse model of HSAN V(Yang et al., 2018). We showed that unlike the homozygotes (NGF^fln/fln^) that often failed to thrive to adulthood, development of survival of the heterozygotes (NGF^fln/+^) showed no impairment. However, they developed a progressive degeneration of small sensory fibers, which could be observed both behaviorally and functionally. Interestingly, the heterozygotes (NGF^fln/+^) showed no obvious structural alterations in the brain including NGF-dependent BFCNs. Accordingly, they did not develop appreciable deficits in learning and memory. These observations with carriers of the mutant allele are in line with data from limited human studies(Minde et al., 2009; Minde et al., 2004b; Minde, 2006). Mechanistically, we showed the mutation resulted in reduced secretion of mature NGF in primary mouse embryonic fibroblasts cultured from the mouse model. We believe these findings will reignite efforts of using NGF as a therapeutic agent, and further would allow to develop ‘painless NGF’ therapy (Capsoni et al., 2011; Cattaneo and Capsoni, 2019; Sung et al., 2019).

## Materials and Methods

### Ethical statement

All experiments involving the use of laboratory animals have been approved by the Institutional Animal Care and Use Committee of University of California San Diego. Surgical and animal procedures were carried out strictly following the NIH Guide for the Care and Use of Laboratory Animals. When possible, we made every effort in our experiments to measure differences between male and female animals.

### Generation of a NGF^R100W^ knockin mouse model

NGF^R100W^ knockin mice were generated by the Model Animal Research Center of Nanjing University (Nanjing, Jiangsu, China) as described previously(Yang et al., 2018). Briefly, the arginine (R) at position 100 of mature NGF (position 221 for proNGF) was mutated to tryptophan (W) by gene targeting (Figure 1A). A targeting vector harboring the A-to-T mutation at the 661 position in exon 3 was constructed by bacterial artificial chromosome (BAC) recombineering, and the mutant allele was subsequently introduced into embryonic stem (ES) cells via homologous recombination. The accompanying neomycin resistance gene served as a selection marker for screening for positive clones. Following confirmation of gene-targeted clones by Southern blot, correctly targeted ES cells were microinjected into donor blastocysts that were transferred to recipient C57BL/6 females to produce chimeric mice. Chimeric mice were bred with C57BL/6 mice to generate heterozygous mice. Genotyping identification of NGF^R100W^ knockin mice was carried out by PCR using the primer pairs: 5-GGGGAAGGAGGGAAGACATA-3 for forward primer and 5-GATTCCCTTAGGAAGGTTCTGG-3 for reverse primer. The following amplification protocol (95°C 5 min; 95°C 30s; 58°C 30s; 72°C 45s; 35 cycles; 72°C 5 min; 15°C hold) was used for genotyping; the expected PCR products were marked for NGF^+/+^, NGF^+/fln^ or NGF^fln/fln^ (Figure 1B).

**Fig 1.**
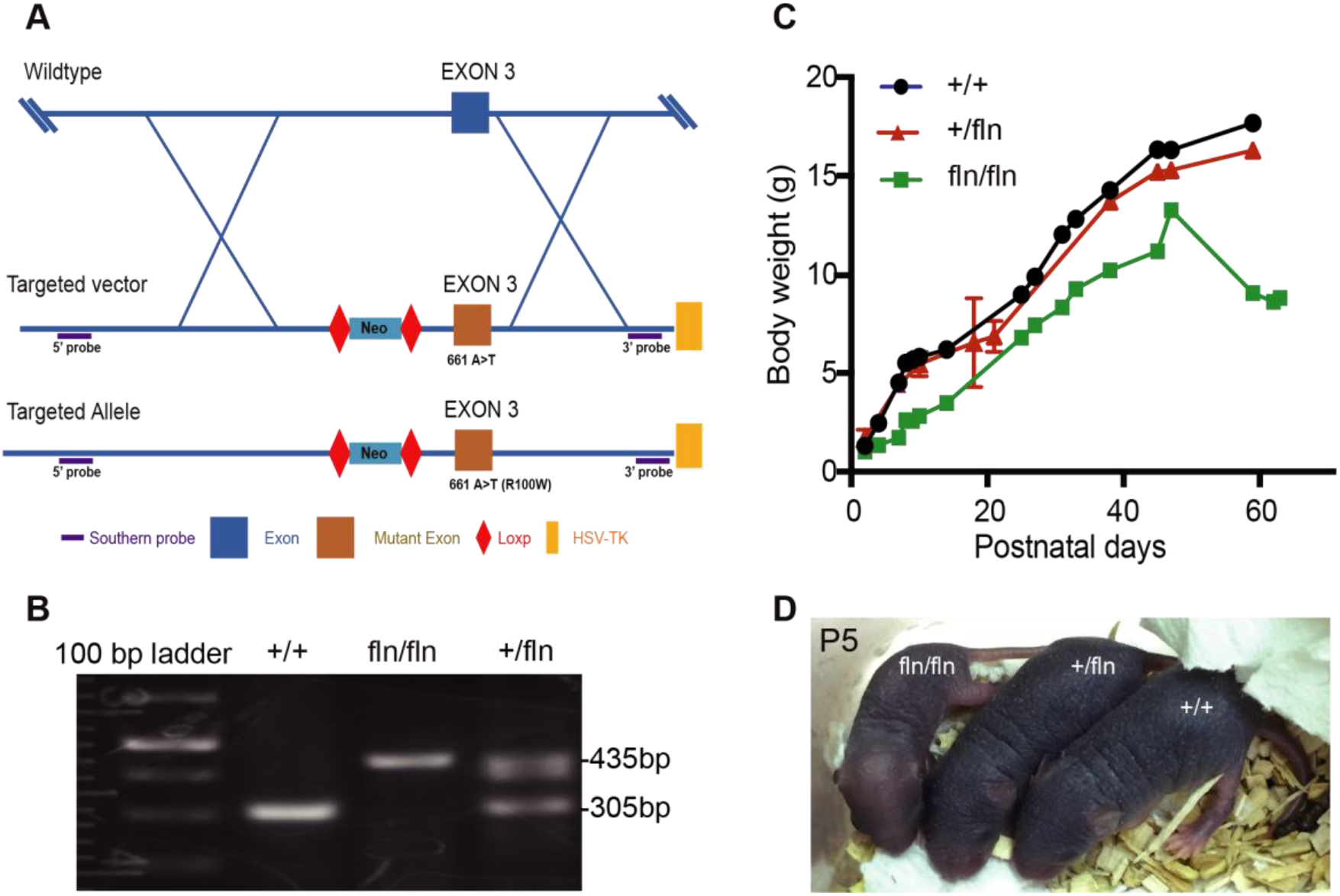
Generation of NGF^R100W^ knockin mice and their body weight phenotypes. (A) NGF^R100W^ knockin mouse model in C57BL/6 background were generated by homologous recombination. (B) Genotyping for wild-type (NGF^+/+^), the heterozygote (NGF^+/fln^) and the homozygote (NGF^fln/fln^). (C) The body weight of three genotype was monitored over a period of 8 weeks (NGF^+/+^, n=3; NGF^+/fln^, n=5; NGF^fln/fln^, n=3). (D) Pups of the three genotypes at postnatal 5 days.

#### Animal housing conditions

All mice were housed in individual cages on a 12/12 h light/dark cycle at 21±2°C. All tests were carried out in a quiet room between 10 AM and 4 PM using sex- and age-matched mice.

### Behavioral tests

#### Nociceptive hot plate test

In nociceptive hot plate test, the mice were placed on a metal hot plate at 55°C (Analgesia Hotplate, Columbus Instruments, Columbus, OH). The latency time to a discomfort reaction (jumping, licking or shaking hind paws) was recorded and immediately remove the mice from the hot plate and return to the cages. The cut-off time was 20 sec.

#### Morris water maze

The Morris water maze was used to evaluate spatial learning and memory as previously described (Spencer et al., 2017). A platform was placed in the center of one quadrant of the pool (diameter 180 cm). Fill the pool with opaque water (24°C) until the platform was submerged 1 cm beneath the water surface. Briefly, the mice were first trained to find a platform with a visible flag on days 1 to 3 and then a submerged hidden platform on days 4 to 7. Each mouse was place into the water facing the wall of the pool at its designated departure point. The departure point was changed randomly between two alternative entry points located at each intersecting line between two zones and same distance from the platform. Each mouse had four consecutive 90 s trials with a brief rest. Mice that failed to reach the platform within 90 s were held by the tail and guided to the platform and stayed on it for another 30 s. On day 8 of probe test, for the first 40 seconds trial, with the removal of the platform, the mice were let to search the pool. Times spent by mice in the target quadrant and # passing of the target zone were recorded. The second 40 seconds trial, the visible platform was placed to the same position and the time to find the platform (latency time) was recorded.

#### Y-Maze spontaneous alternation test

The Y-Maze spontaneous alternation test was performed to measure change in the natural propensity of mice to explore new environments and spatial memory. The test was carried out in a Y-shaped maze with three enclosed black, opaque plastic arms (A, B, C, and 29 cm × 68 cm × 619 cm each) at a 120° angle from each other. The mice were placed into the center of the maze and allowed to freely explore the three arms for 5 minutes. An entry occurs when hind paws of mice were within the arm. The total number of arm entries and the entering sequence were recorded. A spontaneous alternation performance (SAP) occurred when mice entered three successive different arms (e.g. ABC, BCA, CAB, BAC, ACB, CBA). The SAP percentage was calculated with formula: [Total number of SAP/(total arm entries–2)] ×100(Carpenter et al., 2012).

#### Novel object recognition test

A novel object recognition test was performed with a black open-filed chamber (31 × 24 × 20 cm. length: width: height) to assess short-term and long-term memory impairment in mice(Lueptow, 2017). We used three objects that differ in shape and texture: tower of Lego bricks (4 long × 2 wide × 8 cm high), Thermo cell culture polystyrene flask with sand (75ml) and glass Erlenmeyer flask with sand (50ml). Object preference was examined in prior experiments with mice of similar age, and mice have equal preferences for those three objectives. On day one, mice were habituated in the chamber for 10 minutes and return the mouse to its home cage. On day two, two identical objects (tower of Lego bricks) were placed in the chamber, and a mouse was placed to the chamber and allowed to explore both objects for 8 minutes. After the familiarization, it was returned to the home cage. For the short-term memory impairment test, after 30 min interval, one of the familiar towers of Lego bricks used for the memory acquisition was replaced with a new object (cell culture polystyrene flask with sand). The mouse was placed in the chamber for 8 minutes to explore the objects. On day three, for the long-term memory impairment test, after 24 hours interval, one of the towers of Lego bricks was replaced with another new object (glass Erlenmeyer flask with sand) and allowed the mice to explore the objects for 8 minutes. The activities of mice during the test were recorded with a video camera and the time spent exploring each object was assessed. The discrimination ratio was calculated with the formula: discrimination ratio (%)=time spent exploring novel object/total time spent exploring both objects) ×100.

#### Marble burying test

The marble-burying test was performed to evaluate the anxiety-like behavior in mice(Thomas et al., 2009). Twenty marbles were equidistantly placed (5×4) on the bedding surface. After 20 min, the number of marbles buried (>50% of its surface area is covered by bedding) was counted.

#### Light-dark shuttle box test

Light-dark shuttle box test was carried out to evaluate the anxiety-like behavior in mice(Heredia et al., 2014). The apparatus consists (21cm x 42cm x 25cm) of two equally sized chambers, a black chamber and a brightly illuminated white chamber connected by a door. Mice were placed in the dark chamber facing away from the opening and open the door in 3 seconds. Mice were allowed to move freely between the two chambers for 10 min. The latency to first cross to the light chamber and the time spent in the bright chamber were recorded.

#### Social interaction test

Social interaction test was performed to measure the social memory impairment in mice(Kaidanovich-Beilin et al., 2011). The apparatus (40.5cm x 60cm x 22cm) consists of three equally sized chambers connected by doors. Two wire containment cups (a height of 15cm and diameter of 7cm) are placed in the two side chambers respectively and two control mice will be used for each test. Before the test, close the doors and place the mice at the center of the middle chamber for 5min to familiar with its surroundings. After the habituation, for the social affiliation aspect of the test (session I), one of the control mice (Stranger 1) was placed inside one of the cups that were located in one of the side chamber. Another cup was empty. Mouse was placed to the middle chamber and opened the doors to allow free access for the subject mouse to explore each of the three chambers for 10 min. For social novelty/preference session of the test (session II), placed another control mouse (Stranger 2) to the empty cup in the opposite side chamber and monitored the same parameters in session I for another 10 min. The duration of contacts between experimental mouse and empty cup or stranger 1 in session I, or between experimental mouse and stranger 1 or stranger 2 in session II were recorded.

### Immunohistochemistry

#### Histomorphometric analysis of the sciatic nerve

Mice were anesthetized using isoflurane and sacrificed with a guillotine. Sciatic nerves were collected and fixed in 2.5% glutaraldehyde (in 0.1M sodium phosphate buffer) for 24 h at 4℃. Nerves were then rinsed with 0.1M sodium phosphate buffer and postfixed in 2% osmium tetroxide (1ml 4% osmium tetroxide + 1ml 0.2 M sodium phosphate buffer) for 30 min at room temperature. Nerves were rinsed with distilled water twice and then dehydrated through graded ethanol (10 min each in 30%, 50%, 70%, twice in 95.5% for 15 min for each rinse, and twice in 100% for 15 min for each rinse). Samples were rinsed in propylene oxide twice for 15 minutes for each rinse and then put in a mixture of 50% propylene oxide and 50% resin for at least 2 hours. Finally, samples were embedded in 100% resin and sectioned using a glass knife and microtome to 0.5μm semi-thin section. Sections were stained with 2% P-Phenylene Diamine (PPD) at room temperature for approximately 20 minutes and analyzed with a light microscope. The diameter of myelinated fibers was measured using NIH ImageJ.

#### Dorsal root ganglion (DRG) and brain

L3-L5 DRGs were collected and fixed in 4% paraformaldehyde for 30 min at 4°C, and then permeated with 15 % sucrose at 4°C overnight. DRGs were then embedded in Tissue-Tek O.C.T (Electron Microscopy Sciences, Hatfield, PA) and sectioned into 20 μm cryo-sections. Brains were dissected and fixed in 4% paraformaldehyde for 5 days at 4°C. Brains were then sectioned using vibratome to 40μm. Sections were incubated in 1% Triton X, 10% H_2_O_2_ in PBS for 20min at room temperature, washed three times with PBS, blocked 1 h at room temperature with 10% goat serum and then incubated with primary antibodies overnight. Sections were then incubated with biotinylated secondary antibody at 1:100 made in PBS for 30min at room temperature, washed with PBS three times, incubated with avidin-biotin-peroxidase (ABC Kit, VECTOR LABORATORIES, INC. Burlingame, CA) for 1 hour and followed by peroxidase detection (DAB Kit, ABC Kit, VECTOR LABORATORIES, INC. Burlingame, CA). Finally, sections were mounted and analyzed with a light microscope (Leica DMi8 Live Imaging Microscope).

#### Hind paw footpad skin

Skin samples were dissected and fixed in methanol/acetone (1:1) for 30 min at −20°C, rinsed with PBS three times, and incubated in 30% sucrose overnight at 4°C. Tissues were embedded in Tissue-Tek O.C.T (Electron Microscopy Sciences, Hatfield, PA) and sectioned into 45 μm cryo-sections using a Leica Cryostat (Model# CM1900). Sections were incubated in 50mM Glycine for 45min at room temperature, rinsed in PBS three times, blocked in 10% goat serum, 1% BSA, 0.5% Triton X in PBS and then incubated with primary antibodies overnight at 4°C. Primary antibodies were diluted in 10% goat serum, 1% BSA, 0.2% Triton X in PBS. Sections were rinsed three times with PBS and then incubated with secondary antibodies at room temperature for one hour. After washing with PBS three times, sections were mounted and analyzed by confocal microscopy (Leica TCS SP8 Confocal Microscope). Approximately, for samples from each animal, we examined 10-15 sections with 200 μm in length/section. The image stacks were collected at a 0.5 μm step size. Maximal projection images were generated from these stacks and the IENF density was quantified using ImageJ.

### Primary culture of mouse embryonic fibroblasts and ELISA measurement of NGF

Skin samples were dissected from mouse E16-18 (NGF^+/+^, NGF^+/fln^ or NGF^fln/fln^) embryos. Following dissociation, fibroblasts were cultured in 6 well plate in DMEM supplemented with 4.5 g glucose, 10% FBS and 1% penicillin/streptomycin. At 70-80% confluency, the cultures were changed to serum-free medium for 48 hrs. Media were collected and NGF secreted into media was measured by commercial ELISA (Cat. No. OKBB00230, Aviva Systems Biology, San Diego, CA), following manufacture’s protocol.

### Chemicals, antibodies

Unless specified, all chemicals were purchased from Sigma or Fisher. Abs: the UCHL1/PGP9.5 rabbit polyclonal antibody (Cat.No. 14730-1-AP, Proteintech Group, Inc, Rosemont, IL) with a dilution of 1/300 to 1/500. CGRP rabbit antibody (Cat.No 19005, OWL), CGRP mouse antibody (Cat. No 57053, Santa Cruz Technology), anti-TrkA antibody (UCSF?) with dilution of 1/100. NeuN antibody (Cat.No MAB377B, Millipore, Billerica, MA, USA) with dilution of 1/100. Biotinylated horse anti-mouse IgG antibody (Cat.No. BA2000, VECTOR LABORATORIES, INC. Burlingame, CA) with dilution of 1;100. Biotin-conjugated donkey anti-rabbit IgG (Cat.No 711-065-152, Jackson ImmunoResearch, West Grove, PA, USA) with dilution of 1;100.

### Statistical analysis

Statistical analyses were all carried out using GraphPad Prism (GraphPad Software, Inc, La Jolla, CA). For two group comparisons, Student’s t-test was performed. For multiple comparisons, one-way ANOVA or Two-way ANOVA was performed. All data were represented as mean values ± SEM. P<0.05 was considered statistically significant differences.

## RESULTS

### The NGF^R100W^ knockin mouse model

NGF plays a critical role in regulating the differentiation and/or the survival of sensory, sympathetic and BFCN neurons (Casaccia-Bonnefil et al., 1998; Chao, 2003; Cuello et al., 2010; Huang and Reichardt, 2001; Levi-Montalcini, 1987, 2004; Levi-Montalcini et al., 1995; Levi-Montalcini and Hamburger, 1951; Mobley et al., 1986). Although a total knockout of NGF function in mice is postnatally lethal (Crowley et al., 1994), hemizygous NGF mice show only a mild cholinergic deficit with no other apparent phenotypes (Chen et al., 1997). Phenotypic knockout of NGF using antibodies causes a plethora of central and peripheral phenotypes (Ruberti et al., 2000). Studies with NGF^R100W^ have pointed to decoupling of the two important biological functions of NGF: trophic versus nociceptive(Capsoni et al., 2011; Sung et al., 2018; Sung et al., 2019; Yang et al., 2018). We thus generated a NGF^R100W^ knockin mouse model in C57BL/6 background by homologous recombination (Fig 1A, B). The homozygous mice (NGF^fln/fln^) were born with normal body weight but gained weight more slowly after birth with 75% of the mean weights of their wild type mice at 4 weeks of age (Fig 1C). The NGF^fln/fln^ mice also showed developmental delay in many other aspects, including eye opening and hair growth (Fig 1D). Furthermore, observations during the early postnatal days showed that NGF^fln/fln^ mice displayed smaller milk spot in the stomach and often died within the first week of life.

By reducing litter size through moving littermates that has bigger milk spot to a wild-type (NGF^+/+^) surrogate mother, we were able to rescue the NGF^fln/fln^ mice. Some of these mice could ingest more milk and survived up to 6 weeks with one survived for 2 months. These results indicated that NGF^fln/fln^ mice were born live with normal weight but often failed to thrive. The heterozygous mice (NGF^+/fln^), on the other hand, exhibited normal body weight and could not be distinguished from their littermates by their appearance and spontaneous behaviors.

### NGF^fln/fln^ mice exhibits severe loss of PGP9.5-positive intra-epidermal sensory fibers (IENFs) and CGRP-positive small sized neurons in dorsal root ganglia (DRG)

HSAN V patients showed a moderate loss of Aδ fibers and a severe reduction of C fibers (Minde et al., 2004b). Myelinated Aδ and unmyelinated C fibers are the primarily responsible for detecting painful stimuli and transmitting the signal to the brain via the DRG-spinal cord pathway (Dubin and Patapoutian, 2010). Therefore, we first analyzed the density of intra-epidermal sensory fibers (IENFs) in the hind paw skins from postnatal 0 (P0) day mice of all three genotypes. Specific antibody against PGP9.5, a pan-neuronal marker, was used to stain IENFs, including myelinated Aδ fibers and unmyelinated C-fiber (Jolivalt et al., 2016). The average density of IENFs for each genotype was quantified. As illustrated in Fig. 2A and B, The fiber density in the epidermis appeared clearly reduced in NGF^fln/fln^ mice compared with NGF^+/+^ mice (Fig. 2A and B, Movie S1 and S3). Although the NGF^+/fln^ mice at P0 showed a decrease as well, the reduction did not reach significance in comparison with the NGF^+/+^ mice (Fig. 2A and B, Movie S1 and S2).

**Fig 2.**
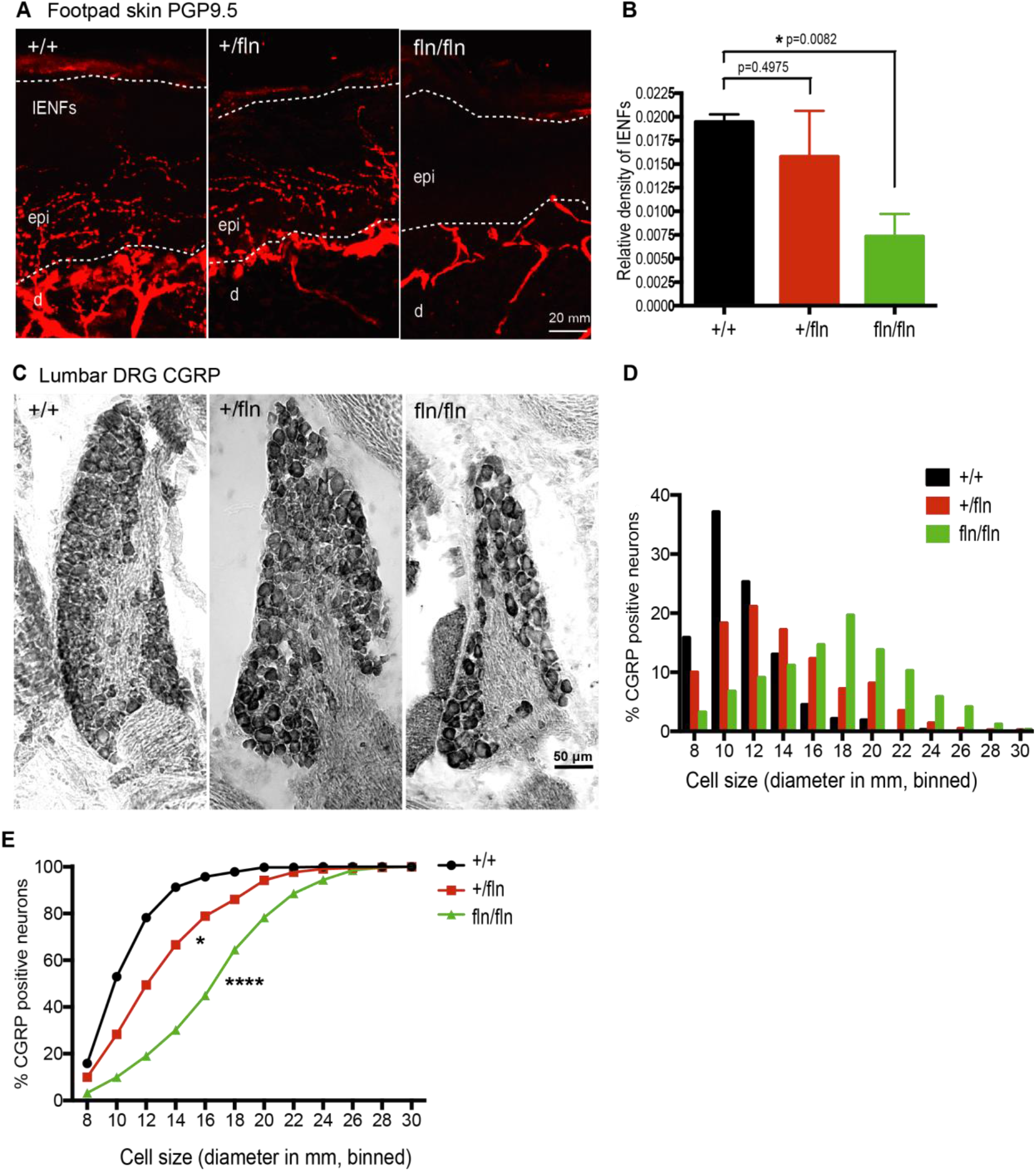
NGF^fln/fln^ mice exhibit loss of PGP9.5-positive intra-epidermal sensory fibers (IENF) and CGRP-positive neurons in DRGs at postnatal 0 day. (A) Sections of footpad skin from P0 mice were stained with antibodies to the pan-nerve fiber marker PGP9.5. (B) Quantification of PGP9.5-positive nerve fibers intensities in epidermis of footpad skin. To estimate the density of IENF in the epidermis, at least 12-15 images of each animal were analyzed throughout the section thickness and entire length of each image. About 2600 μm of footskin per each animal was analyzed. epi: epidermis; d: dermis. Mean±SEM, * = p<0.05 by unpaired t test. NGF^+/+^, n=3; NGF^+/fln^, n=3; NGF^fln/fln^, n=3. (C) Immunoreactivity to CGRP in representative sections of lumbar DRG from P0 mice. (D) Cell size distribution of CGRP-positive neurons in DRG. NGF^+/+^, n=3 (423 cells); NGF^+/fln^, n=3 (431cells); NGF^fln/fln^, n=3 (341 cells). (E) Cumulative distribution of CGRP-positive neurons in DRG. *=p<0.05, ****=p<0.0001, one-way ANOVA test followed by Dunn’s multiple comparisons test (compared with +/+ group).

DRG neurons convey sensory information from the periphery to the CNS (Meyer RA, 2006; Willis WD, 2004). Pain sensation is primarily conveyed by small DRG neurons (≤ 30 um)(Li et al., 2016; Meyer RA, 2006; Willis WD, 2004), which are predominantly marked by the calcitonin gene-related peptide (CGRP) (Li et al., 2016). To investigate if this population of DRG neurons were affected by the NGF^R100W^ mutant allele(s), we extracted DRGs from NGF^+/+^, NGF^+/fln^, and NGF^fln/fln^ P0 pups. DRGs were processed and stained with a specific antibody against CGRP. Representative images of the three genotypes are presented in Fig 2C. Strikingly, the CGRP-positive neurons in the NGF^fln/fln^ samples showed a significant reduction in the small size neurons (Fig 2C). The soma sizes of CGRP-positive neurons were quantified and the size distributions for each genotype are shown in histograms and cumulative percentage plot (Fig 2D and E). The majority of CGRP-positive neurons (∼90%) in the NGF^+/+^ samples fell within the range of 8-14 μm in diameter. DRGs from NGF^+/fln^ exhibited a similar size distribution pattern, although with a slight increase in the range of 18-22 μm. The NGF^fln/fln^ sample showed a marked decrease to ∼35% percentile in small size neurons (8-14 μm) as compared to ∼90 percentile in NGF^+/+^ (Fig 2D), while the large size neurons (≥16 μm) in NGF^fln/fln^ was increased to ∼65 percentile from ∼10 percentile in the NGF^+/+^ DRGs. The is more evident in the cumulative percentage plot that was significantly shifted to the right in NGF^fln/fln^ as compared with NGF^+/+^. Taken together, these data have demonstrated that the NGF^R100W^ mutant allele, especially presented in two copies, results in a marked reduction in the number of both IENFs in the skin and CGRP-positive small size neurons in DRGs.

### NGF^fln/fln^ mice show severe deficits in sensing hot temperature

Given that both IENFs and small size DRG neurons were severely reduced in the NGF^fln/fln^ mice, accordingly we predicted that these mice would develop severe deficts in perceiving painful stimuli. We next used a hot plate assay (55°C) to test the threshold of 2 month old mice to thermal stimuli using a 20 sec cutoff. As showed in Fig.3 A, there was no significant difference between the NGF^+/+^ and the NGF^+/fln^ mice with an average latency time of ∼11 sec for both genotypes. However, the latency time for the NGF^fln/fln^ mice was significantly increased as compared to either the NGF^+/+^ or the NGF^+/fln^ mice. Strinkingly, the NGF^fln/fln^ mice were totally insenitive to the 55°C thermal stimuli and showed no signs of discomfort on the hot plate. In the end, they had to be immediately removed at the 20s cutoff. Together with the observation of severe reduction of both IENFs and small size DRG neurons in the NGF^fln/fln^ P0 pups, these results provide strong evidence that the NGF^fln/fln^ mice have lost the ability to perceive and respond to noxious stimuli, a telltale sign for severe peripheral sensory neuropathy. Therefore, the NGF^R100W^ knockin mouse model recapitulates one of the most prominent clincial manifestations of peripheral sensory neuropathy in HSAN V patients (Einarsdottir et al., 2004; Minde et al., 2009; Minde et al., 2006; Minde et al., 2004b; Minde, 2006).

**Fig 3.**
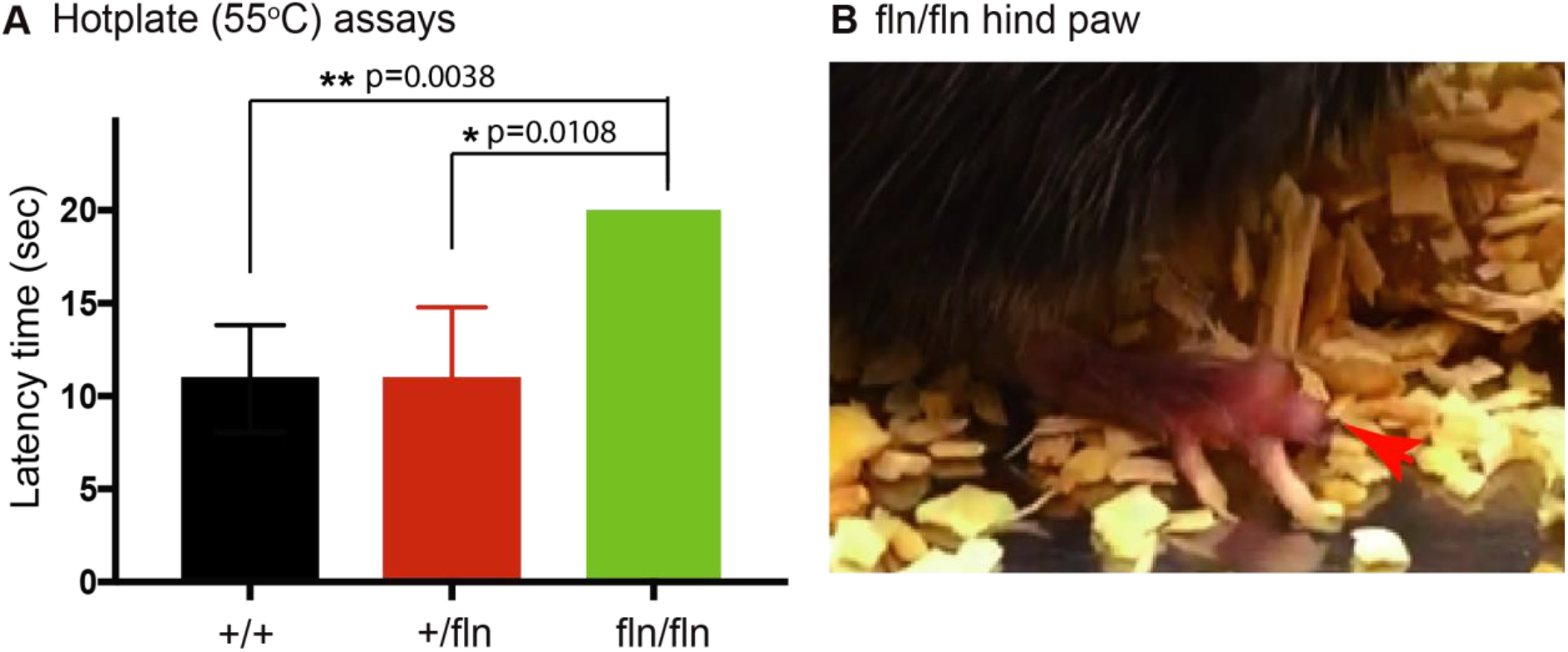
NGF^fln/fln^ display decreased responsiveness to pain at 2 month. (A) The NGF^fln/fln^, not NGF^+/fln^, mice showed significant deficits (20 sec, cutoff time) in 55°C hot plate assays at 2 month. Mean±SEM, * = p<0.05, ** = p<0.01 by unpaired t test. NGF^+/+^, n=39; NGF^+/fln^, n=29; NGF^fln/fln^, n=2. (B) NGF^fln/fln^ showed serious hind paw damage after hot plate test.

### NGF^-/fln^ mice show loss of small size CGRP-positive neurons in DRG with age

Although HSAN V is classified as an autosomal recessive disease (Einarsdottir et al., 2004), heterozygous patients have been also shown to develop sensory neuropathy albeit to a lesser severity with an adult onset (Minde et al., 2009; Minde et al., 2004b; Perini et al., 2016; Sagafos et al., 2016). Since the NGF^fln/fln^ mice often failed to survive to adulthood in sufficient numbers that made it difficult for detailed, systematic and in-depth analysis, we decided to focus our effort on the NGF^+/fln^ mice herein.

We measured the density of CGRP- and TrkA-positive neurons in DRGs in the NGF^+/fln^ mice at different ages (4- and 10-months) and compared with their littermate controls. Representative images along with qunatitative results are shown in Fig 4A and B. Similarly to the results for IENFs, the NGF^+/fln^ mice at 4 months of age did not show any differenece from their same age NGF^+/+^ littermates with regarding the number of CGRP-positive, TrkA-positive or CGRP/TrkA-positive neurons (Fig 4A and B). However, these vaules showed marked changes at 10 months in NGF^+/fln^ mice compared to NGF^+/+^: the percentile of small size CGRP-positive and CGRP/TrkA-positive neurons were reduced, while the percentile of TrkA-positive fibers was not significantly affected (Fig 4A and B).

**Fig 4.**
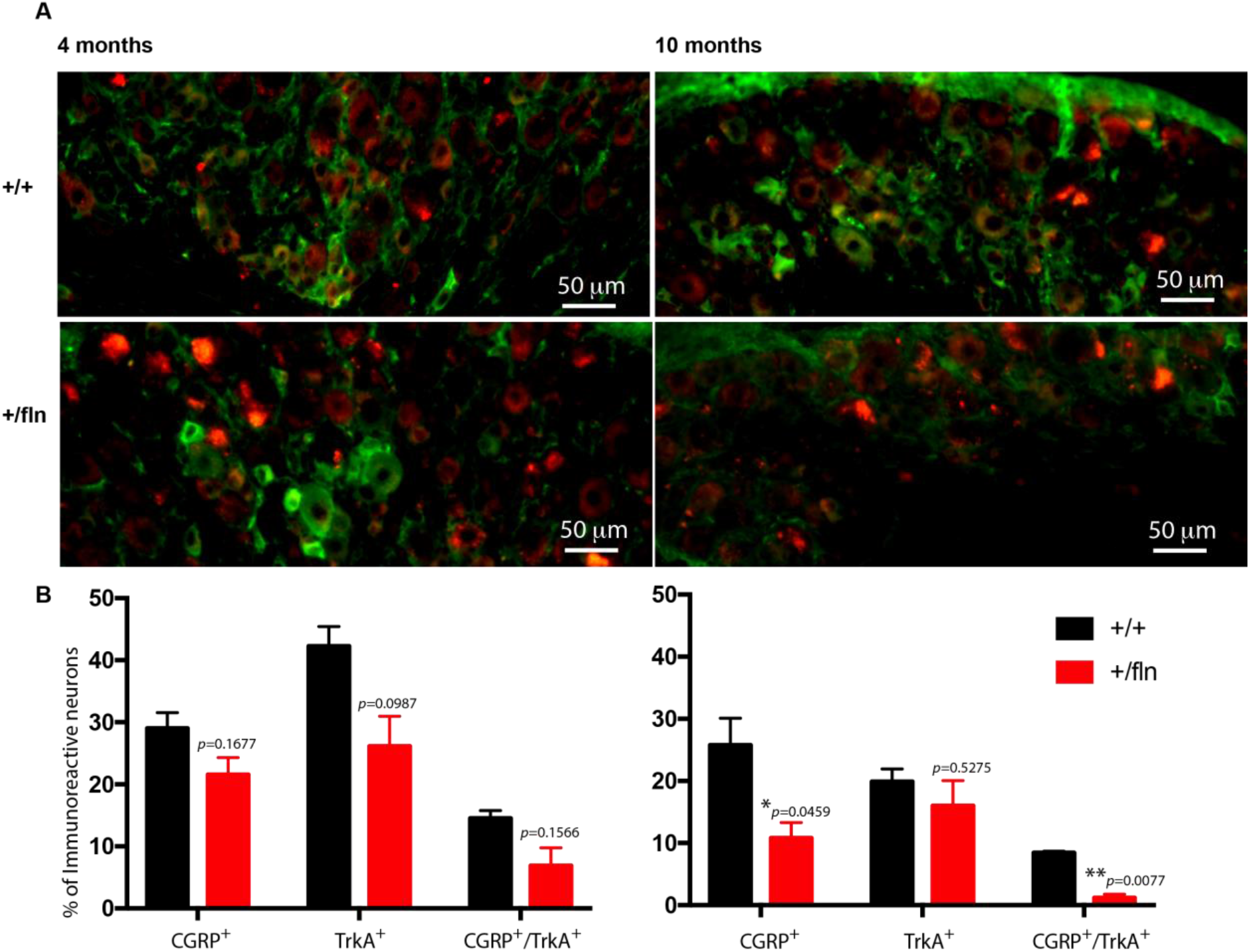
NGF^+/fln^ mice exhibit loss of CGRP-positive small sized neurons in DRG with age. (A), (B), (C) Immunoreactivity to CGRP in representative sections of lumbar DRG from 2, 9 and 18 month NGF^+/+^ and NGF^+/fln^ mice respectively. (E), (F), (G) Cell size distribution of CGRP-positive neurons in DRG from 2, 9 and 18 month NGF^+/+^ and NGF^+/fln^ mice respectively. (H), (I), (J) cumulative distribution of CGRP-positive neurons in DRG from 2, 9 and 18 month NGF^+/+^ and NGF^+/fln^ mice respectively. One pair mice per group, 2 month (NGF^+/+^, 367 cells; NGF^+/fln^ 333 cells), 9 month (NGF^+/+^, 413 cells; NGF^+/fln^, 472 cells), 18 month (NGF^+/+^, 407cells; NGF^+/fln^, 301 cells), ****=p<0.0001 by paired t test.

Since heterozygous patients for NGF^R100W^ showed a moderate loss of Aδ fiber in sural nerve (Minde et al., 2004a), we extracted sciatic nerve and stained myelinated fibers with p-phenylene diamine (PPD) at different ages (2-, 9- and 18-months). Our results have revealed that the median values of myelinated fiber diameters in both NGF^+/+^ and NGF^+/fln^ mice at 2-, 9- and 18 months of age were between 6 and 12μm (Fig 5 A-C). Statistical analysis showed non-significant difference between the two genotypes in size distribution of myelinated fiber diameter. Taken together, these results have demonstrated that a single NGF^R100W^ mutant allele results in a progressive loss of CGRP-positive small sized sensory neurons in DRGs with no apparent loss or shrinkage of myelinated fibers in their sciatic nerve fibers.

**Fig 5.**
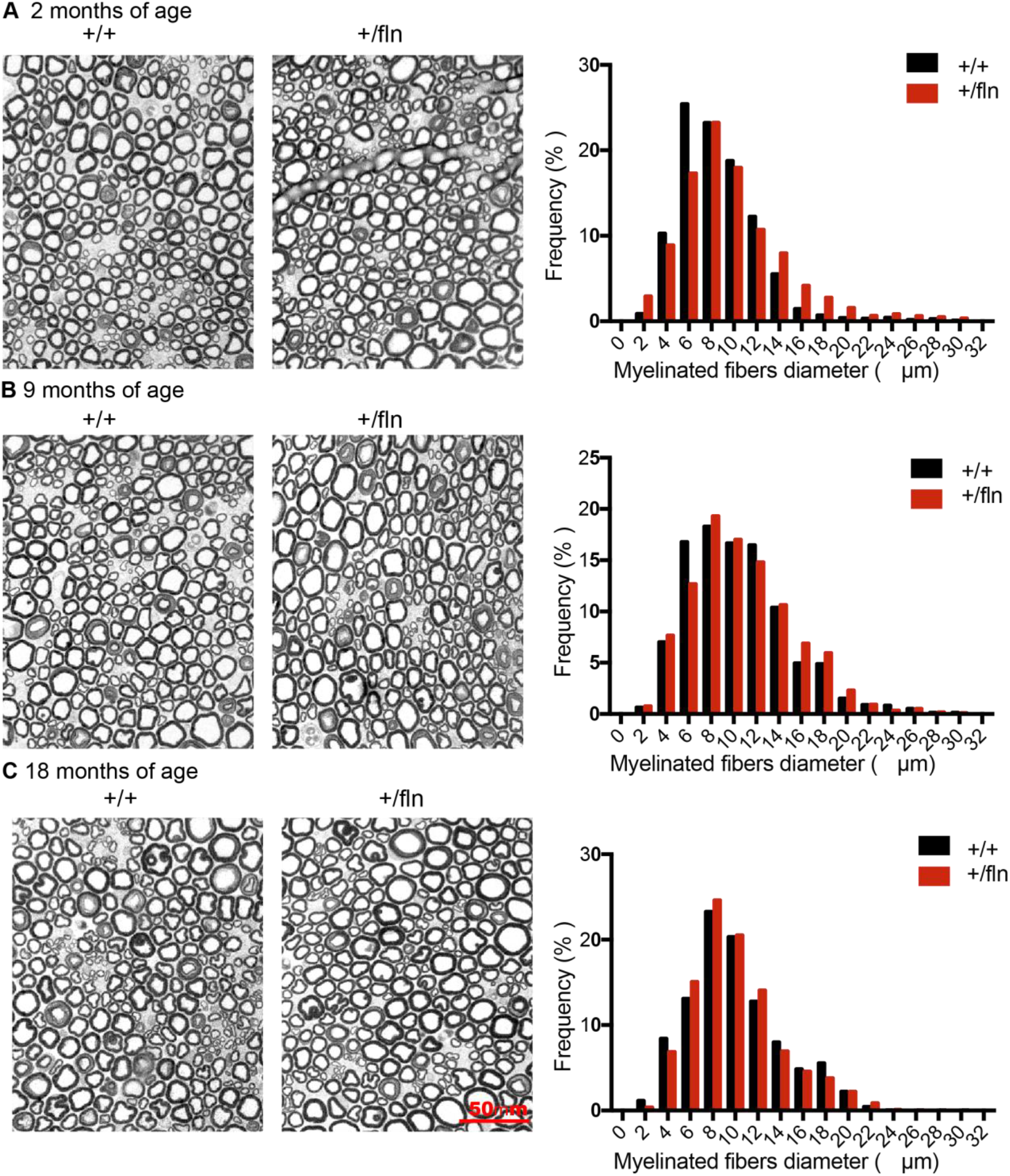
*P*-Phenylene Diamine (PPD) staining of the myelinated fibers in sciatic nerve. (A), (B), Representative sections of the myelinated fibers in sciatic nerve from 2, 9 and 18 month NGF^+/+^ and NGF^+/fln^ mice respectively. Side panel was the size distribution of the myelinated fibers. One pair mice per group, 2 month (NGF^+/+^, 2718 fibers; NGF^+/fln^ 3682 fibers), 9 month (NGF^+/+^, 1685 fibers; NGF^+/fln^, 3524 fibers), 18 month (NGF^+/+^, 1661 fibers; NGF^+/fln^, 806 fibers).

### NGF^+/fln^ mice exhibit progressive degeneration of PGP9.5-positive IENFs

At P0, NGF^+/fln^ mice showed no significant reduction in the density of IENFs in the skin as compared to their NGF^+/+^ littermates (Fig 2). More importantly, the NGF^+/fln^ mice showed no sensory deficits in hot plate assays at 2 months of age (Fig 3), suggesting that they developed normally to adulthood. To investigate if NGF^+/fln^ mice developed progressive sensory neuropathy, as observed in HSAN V heterozygous patients(Minde et al., 2004a), we next examined the NGF^+/fln^ mice at different ages (2, 9, 18 months) using their NGF^+/+^ littermates as controls. Using the same methods as in Figure 3, we quantified the density of IENFs in the hind paw skin. Representative images of skin sections stained with anti-PGP9.5 antibodies are shown in Fig.6 A. Our results have revealed that the NGF^+/fln^ mice indeed showed a progressive loss of IENFs. Quantitative analysis showed no signifciant differences between NGF^+/fln^ and NGF^+/+^ mice at 2 months of age (Fig 6 B). At 9 months of age, the NGF^+/fln^ mice showed marked reduction in the IENF density in the skin (Fig 6 A, B). At 18 months of age, IENFs were hardly detected in the NGF^+/fln^ mice (Fig 6 A and B, Movie S4 and S5). Taken together, these results have demonstrated that a single NGF^R100W^ mutant allele results in a progressive degenerative phenotype of small sensory neurons. These findings are consistent with clinical manifestations of HSAN V patients.

**Fig 6.**
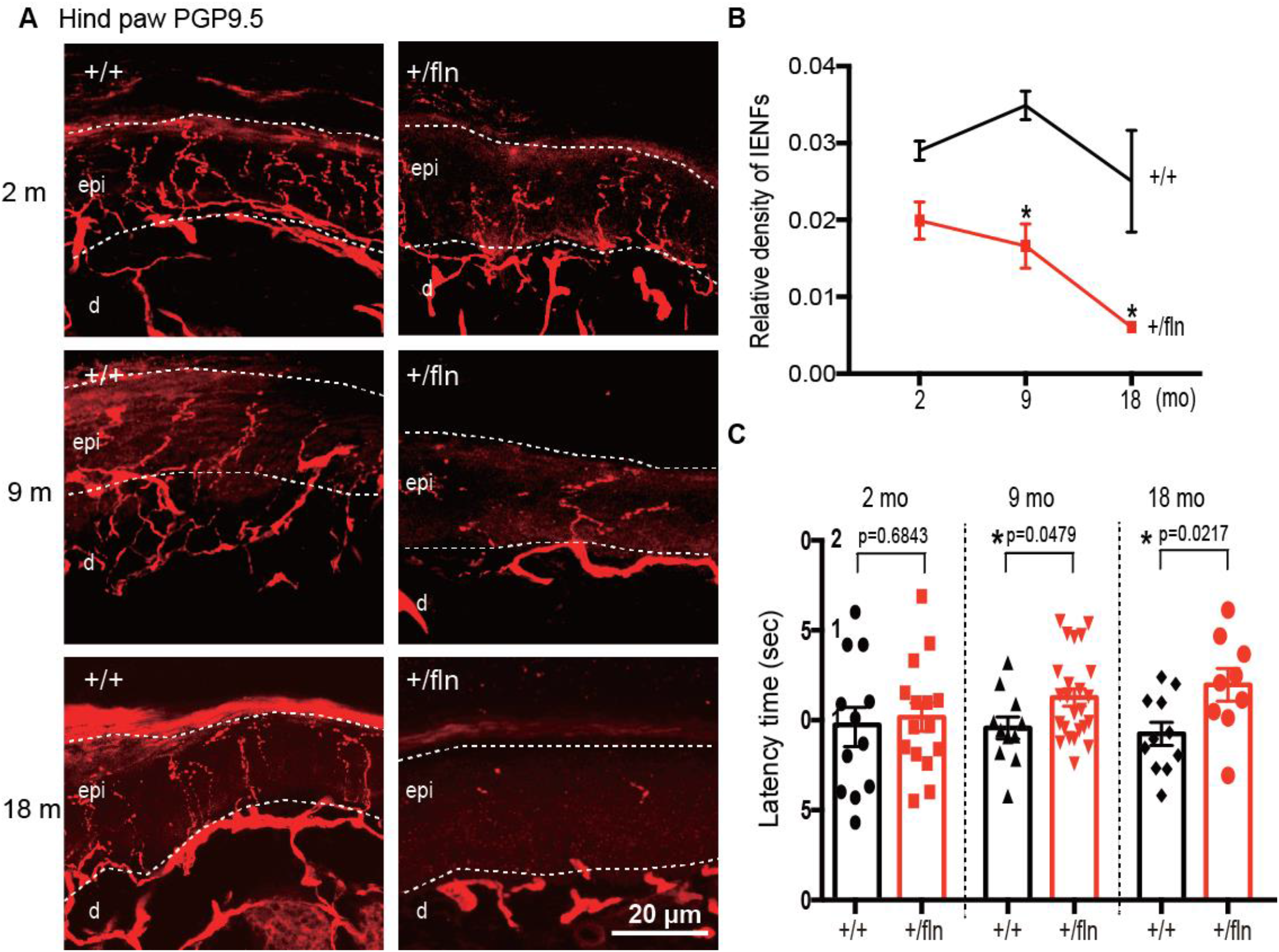
NGF^+/fln^ mice exhibit PGP9.5-positive intra-epidermal sensory fibers (IENFs) degeneration with age and develop progressive sensory deficits. (A) Sections of footpad skin from 2, 9 and 18 month old mice were stained with antibodies to the pan-nerve fiber marker PGP9.5. (B) Quantification of PGP9.5-positive nerve fibers intensities in epidermis of footpad skin. To estimate the density of IENF in the epidermis, at least 12-15 images of each animal were analyzed throughout the section thickness and entire length of each image. About 2600 um of footskin per each animal was analyzed. epi: epidermis; d: dermis. Mean±SEM, * = p<0.05 by two-way ANOVA test. NGF^+/+^, n=3; NGF^+/fln^, n=3; NGF^fln/fln^, n=3. (C) Hot plate assays at 2, 9 and 18 month. Mean±SEM, * = p<0.05 by two-way ANOVA. 2 month (NGF^+/+^, n=12; NGF^+/fln^, n=15), 9 month (NGF^+/+^, n=11; NGF^+/fln^, n=27), 18 month (NGF^+/+^, n=11; NGF^+/fln^, n=9).

### NGF^+/fln^ mice develop progressive nociceptive deficits

Given that NGF^+/fln^ mice showed progressive structural degeneration: reduction in both the numbers of IENFs and small sensory DRG neurons, we predicted that these mice would develop nociceptive deficits. We again used the hot plate assay to assess the pain sensitivity of NGF^+/fln^ mice of 2-, 9- and 18-months of age in comparison with their respective NGF^+/+^ littermate controls. Consistent with the results from studies of IENFs, DRGs, our results demonstrate that the NGF^+/fln^ mice began to show deficits at 9 months of age with the deficits becoming more pronounced at 18 month (Fig. 6 C). These results indicate that NGF^+/fln^ mice develop progressive nociceptive deficits.

### NGF^+/fln^ mice exhibit no marked structural changes in the brain

The impact of NGF^R100W^ mutation on progressive degeneration of the PNS both structurally and functionally raised the question if the CNS was also affected in our mouse model of HSAN V. NGF is mainly produced in the hippocampus and retrogradely transported to the basal forebrain to support survival and differentiation of BFCNs (Mobley et al., 1986; Pezet and McMahon, 2006). To address this, we examined the brain sections from 12-month-old NGF^+/fln^ and NGF^+/+^ mice. Neurons were identified by immunostaining the sections with a specific antibody to NeuN, a marker of neurons. We were particularly focused on the hippocampus and basal forebrain. As illustrated in Fig 7 A, NeuN-immunoreactive neurons were densely compacted in the stratum pyramidale of hippocampal CA1 and CA3 regions and in the dentate gyrus (DG) regions in both the NGF^+/fln^ and NGF^+/+^ brains. We measured the thickness of CA1, which revealed not significant difference between NGF^+/+^ (34.48±0.65 μm) and _NGF^+/fln^ (35.52±1.01 μm) mice hippocampus. (Fig 7 A and C). To determine the impact of NGF^R100W^ in the development of basal forebrain, we analyzed the density of medial septum by quantifying the average NeuN-positive fluorescence intensity in the medial septum. Our results showed that the density of NeuN-immunoreactive neurons was not significantly different between 12-month-old NGF^+/+^ (30804±2367 A.U./mm^2^) and NGF^+/fln^ (33021±8066 A.U./mm^2^) mice (Fig 7 B and C).

**Fig 7.**
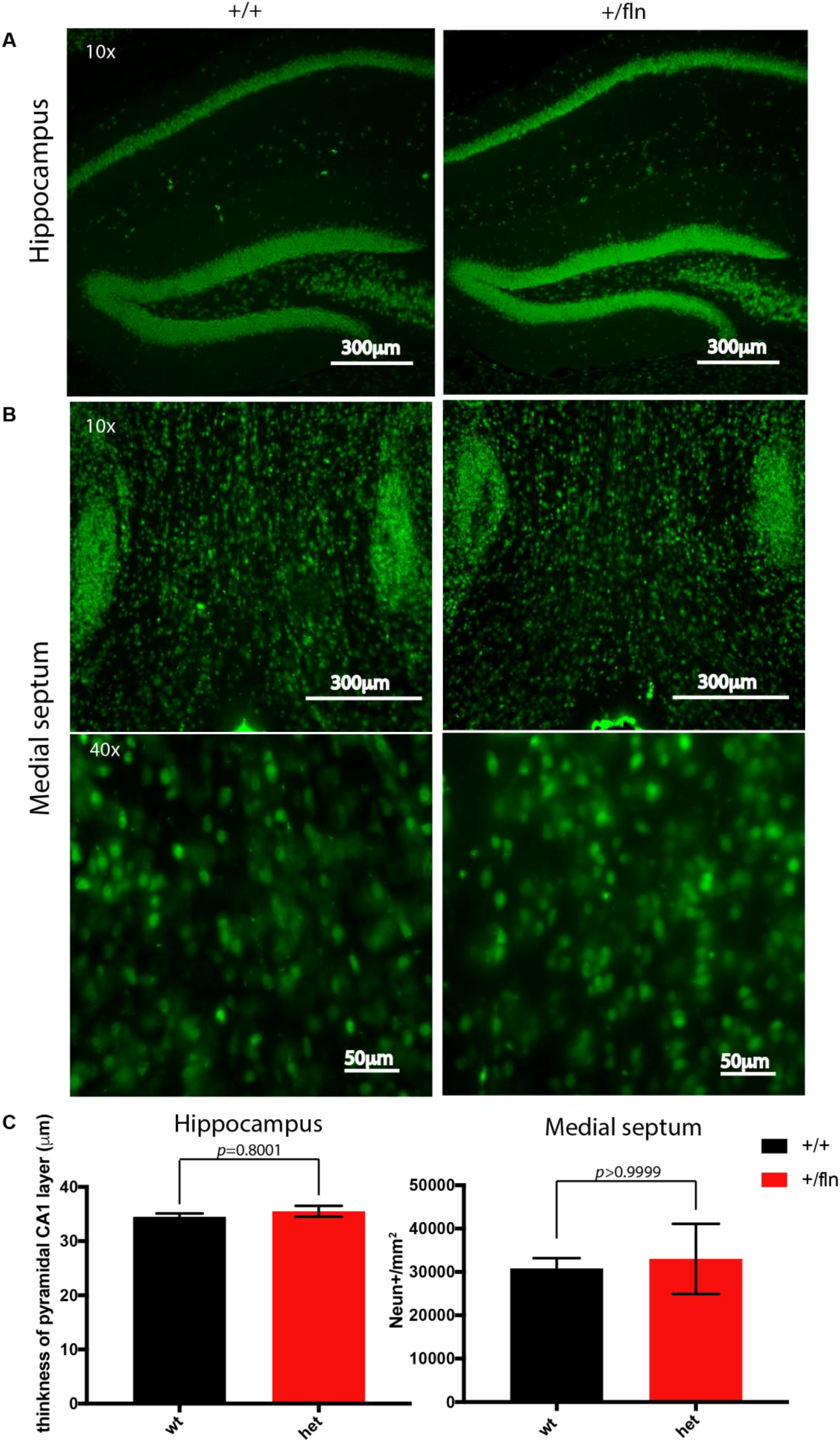
Comparison of the cytoarchitecture of the hippocampus and basal forebrain using NeuN immunostaining between 12-month-old NGF^+/fln^ and NGF^+/+^ mice. (A) Representative images of NeuN-positive cells in CA1 and DG regions of the hippocampus from 12-month-old NGF^+/+^ and NGF^+/fln^ mice. CA1, cornu ammonis 1;; DG, dentate gyrus. (B) Representative images of NeuN-positive cells in medial septum of basal forebrain from 12-month-old NGF^+/+^ and NGF^+/fln^ mice. MS, medial septum. NGF^+/+^, n=3; NGF^+/fln^, n=3. (C) Quantification of NeuN-positive staining in CA1 layer and medial septum.

In the central nerve system, sensations such as pressure or pinprick are registered in the somatosensory cortex, pain is perceived in anterior cingulate cortex (ACC) where emotional reaction to pain is dealt with concomitantly(Price, 2000). It has been suggested that compared to healthy individuals, NGF^R100W^ carriers seem to show weaker blood-oxygen-level-dependent response in their ACC regions using functional MRI (Morrison et al., 2011). Given that pain perception is significantly reduced in NGF^+/fln^ at the age of 9 months, we investigated if the progressive loss of pain perception involved any central component such as changes in ACC. We quantified NeuN-positive neurons in ACC, as marked by the red dashed box in the sagittal brain section of NGF^+/+^ and NGF^+/fln^ mice (Fig 8A). Our data show there was no significant difference in the density of NeuN-positive signals between NGF^+/+^ (1903±88.96 A.U./mm^2^) and NGF^+/fln^ (1689±65.52 A.U./mm^2^) (Fig 8B and C). Taken together, these results have indicated that NGF^+/fln^ mice show no apparent structural changes in their ACC that could also contribute to loss of pain perception.

**Fig 8.**
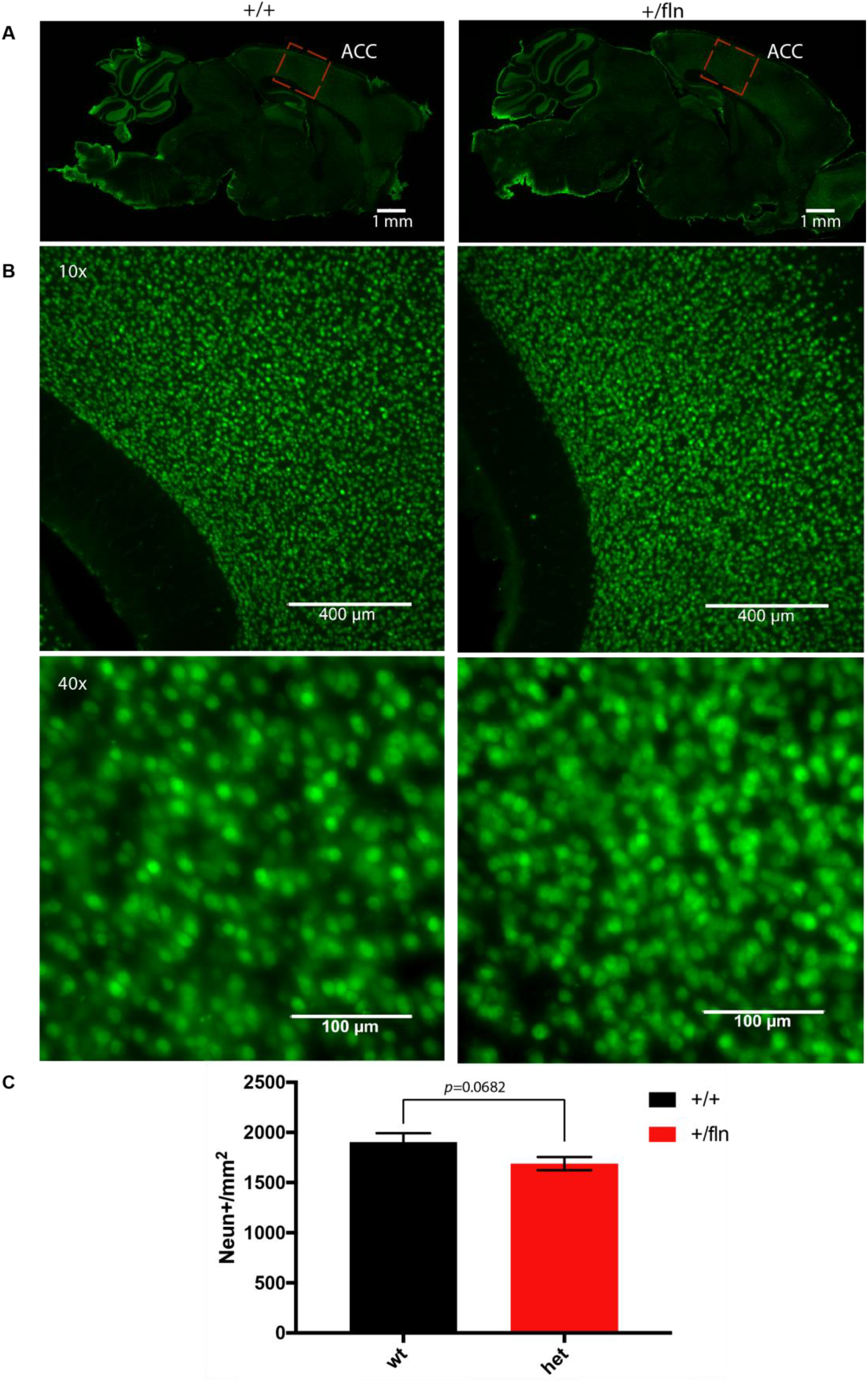
Comparison of the cell density of ACC through NeuN immunostaining between 12-month-old NGF^+/fln^ and NGF^+/+^ mice. (A) Representative images of sagittal section of mice brain. Images were built after stitching and combining multiple 10X microscopic NeuN immunoreactive images. (B) Representative images of NeuN positive cells in ACC from 12-month-old NGF^+/+^ and NGF^+/fln^ mice. NGF^+/+^, n=3; NGF^+/fln^, n=3 (C) Quantification of NeuN-positive staining in ACC.

### NGF^+/fln^ mice show no apparent deficits in CNS dysfunction

One of the major clinical features that distinguish the NGF^R100W^-associated HSAN V patients from HSAN IV associated with mutations in TrkA is that HSAN V patients do not suffer from overt cognitive deficits (Capsoni, 2014; de Andrade et al., 2008; Einarsdottir et al., 2004; Haga et al., 2015; Minde et al., 2004b). To test if this is the case in our HSAN V mouse model, we performed a battery of behavioral tests to assess cognitive functions.

The Y-Maze spontaneous alternation test was performed to evaluate the working memory(Carpenter et al., 2012). Mice prefer to enter a new arm of the maze rather than returning to one that was previously explored and recall the order of the arm entry. As illustrated in Fig 9A, there were no significant differences in the number of arm entries or alternation rate between NGF^+/+^ and NGF^+/fln^ mice at either 9- or 18 months of age. This data indicated that there was no working memory impairment in NGF^+/fln^ mice.

**Fig 9.**
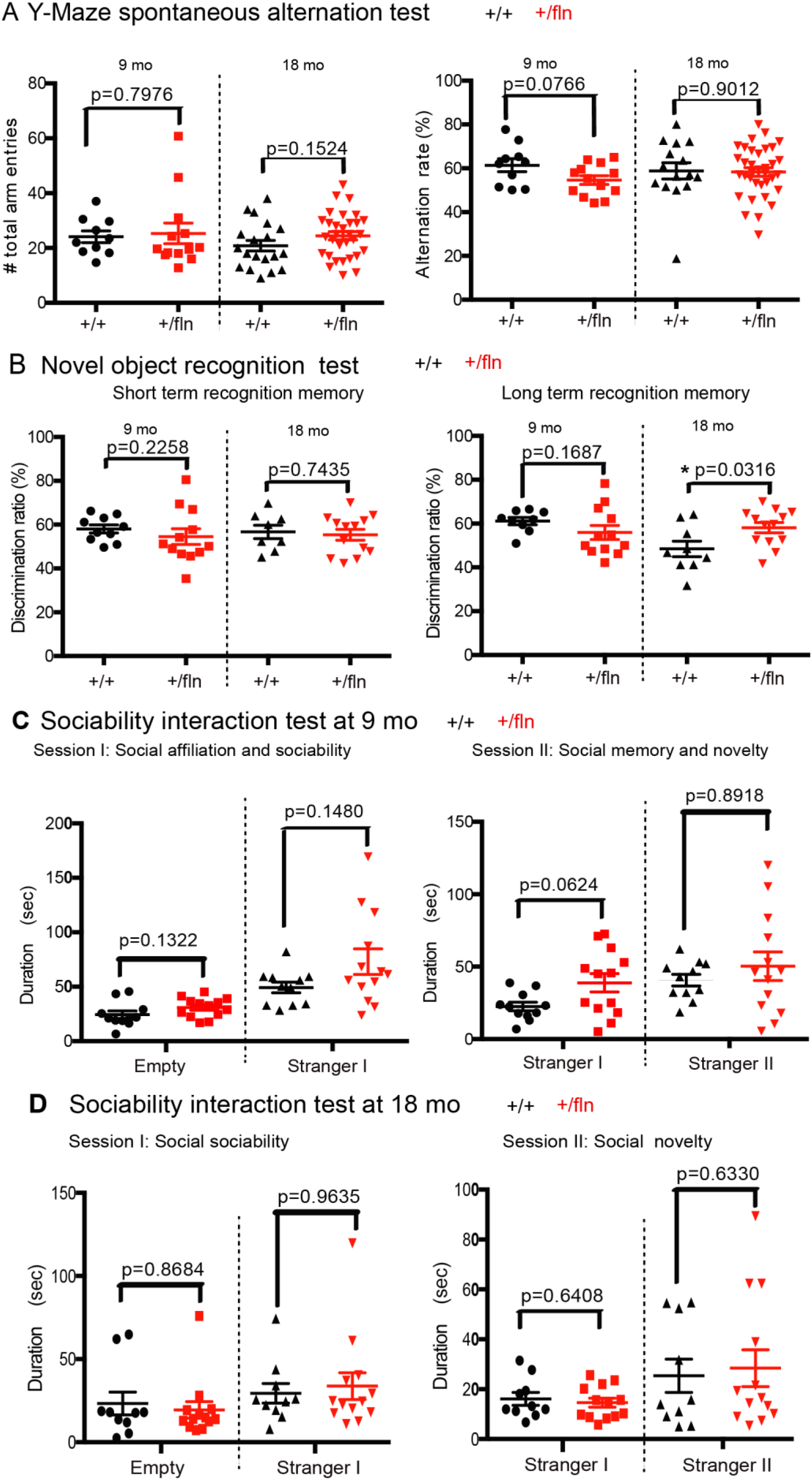
Behavioral analysis of cognitive functions in 9 and 18 month old mice. (A) Y-Maze spontaneous alternation test. Alternation rate (%)=[Total number of SAP/(total arm entries–2)] ×100. Mean ± s.e.m, by unpaired t test. 9 month (NGF^+/+^, n=10; NGF^+/fln^, n=13), 18 month (NGF^+/+^, n=15; NGF^+/fln^, n=35). (B) Novel object recognition test. Discrimination ratio (%)=time spent exploring novel object/total time spent exploring both objects) ×100. Mean ± SEM, * = p<0.05 by unpaired t test. 9 month (NGF^+/+^, n=9; NGF^+/fln^, n=12), 18 month (NGF^+/+^, n=9; NGF^+/fln^, n=13). (C) (D) Sociability interaction Test. Mean±SEM, by unpaired t test. 9 month (NGF^+/+^, n=11; NGF^+/fln^, n=13), 18 month (NGF^+/+^, n=10; NGF^+/fln^, n=13).

We next used a novel object recognition assay to assess short- and long-term memory (Lueptow, 2017). Following acclimating to a designated object (this will serve as the familiar object), the test subject was given the familiar object and a novel object at either 30 min or 24 hours for measuring short- and long-term memory, respectively. We found that at 9 months of age, NGF^+/fln^ mice displayed no difference from their NGF^+/fln^ littermate controls in either short- or long-term memory (Fig 9B).

We then tested these mice for social interactions, which are critical for mice to maintain social hierarchy and make mate choices (Kaidanovich-Beilin et al., 2011). The three-chamber sociability interaction test was performed to measure both the sociability and social novelty. In Session I of this test, a test mouse normally prefers to interact with another mouse over empty cup, the time that this test subject spent with the live mouse is designated sociability. In Session II of the test, the test subject will be provided with either the familiar mouse or another novel mouse, the time that the test mouse spent with the novel mouse is designated as social novelty. As shown in Fig 9C and D, there were no significant differences between NGF^+/fln^ mice at either 9- or 18-months of age and their age-matched littermate NGF^+/+^ controls. We conclude that NGF^+/fln^ mice show no apparent impairment in either their sociability or social novelty.

We also used the marble-burying test to investigate whether NGF^+/fln^ mice displayed anxiety-like behavior at 9 months old(Thomas et al., 2009). When placed in a cage with marbles, mice exhibit digging behavior and bury the marbles, which is seen as anxiety related. As showed in Fig 10A, although not statistically significant difference, there was a trend that NGF^+/fln^ mice buried more marbles than NGF^+/+^ mice indicating NGF^+/fln^ mice werelikely more anxious. To further confirm this, we next performed dark-light shuttle box test using 9- and 18-month old mice, which is also extensively used to test anxiety-like behavior (Heredia et al., 2014). Typically, mice have natural spontaneous exploratory behaviors when placed in a new environment and an innate aversion to brightly illuminated areas. When the test subject was placed in the dark-light shuttle box, its exploratory activity will reflect the balance between those two tendencies and will be influenced by the level of anxiety. Decreased time spent in the light chamber has been used as an index of anxiety. Our results showed that NGF^+/fln^ mice at 9 and 18-months of age displayed no significant difference from age-matched littermate NGF^+/+^ controls in terms of either the time exiting the dark chamber or the time spent in the light chamber (Fig 10B). These results suggest that NGF^+/fln^ mice have no anxiety-like behavior.

**Fig 10.**
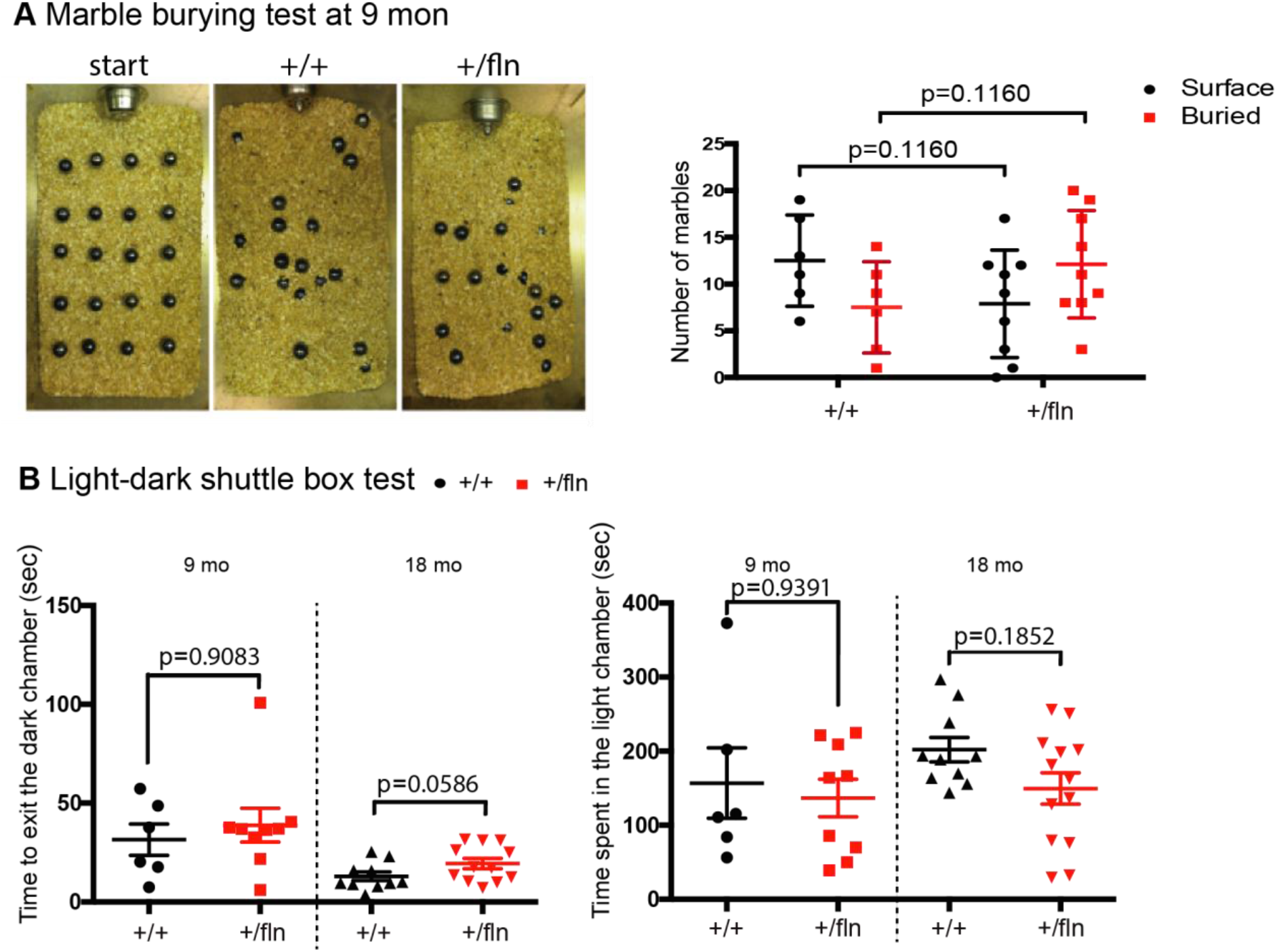
Anxiety-like behavioral test. (A) Marble burying test at 9 month. Mean±SEM, by unpaired t test. NGF^+/+^, n=9, NGF^+/fln^, n=18. (B) Light-dark shuttle box test at 9 and 18 month. Mean±SEM, by unpaired t test. 9 month (NGF^+/+^, n=6; NGF^+/fln^, n=9), 18 month (NGF^+/+^, n=10; NGF^+/fln^, n=13).

To test if NGF^+/fln^ mice at 18 month showed any deficit in their executive function, we used the Morris water maze (Fig 11A) to evaluate spatial learning and memory (Spencer et al., 2017). Test mice were first trained to find a platform with a visible flag on days 1 to 3 and then a submerged hidden platform on days 4 to 7. The path length and time to find the platform (latency time) was recorded. Our result showed the NGF^+/fln^ mice did not differ significantly from NGF^+/+^ mice in training curve (Fig 11B-C). On day 8 of the probe test, for the first 40 seconds trial (session I) with the removal of the platform, mice were let to search in pool. We recorded both the time that mice spent in the target quadrant and the number that mice passed the target zone. Our results showed no difference in either measurements (Fig 11D, E) between NGF^+/fln^ and age-matched littermate NGF^+/+^ controls. In the second 40 seconds trial (session II), the visible platform was re-placed and the latency time was recorded. Again, our test results revealed no significant difference in the latencies between NGF^+/fln^ and NGF^+/+^ mice (Fig 11F).

Taken together, these findings are consistent with clinical findings of HSAN V patients(Capsoni, 2014). NGF^+/fln^ mice show progressive peripheral degeneration both structurally and functionally while developing no marked deficits in their CNS dysfunction.

**Fig 11.**
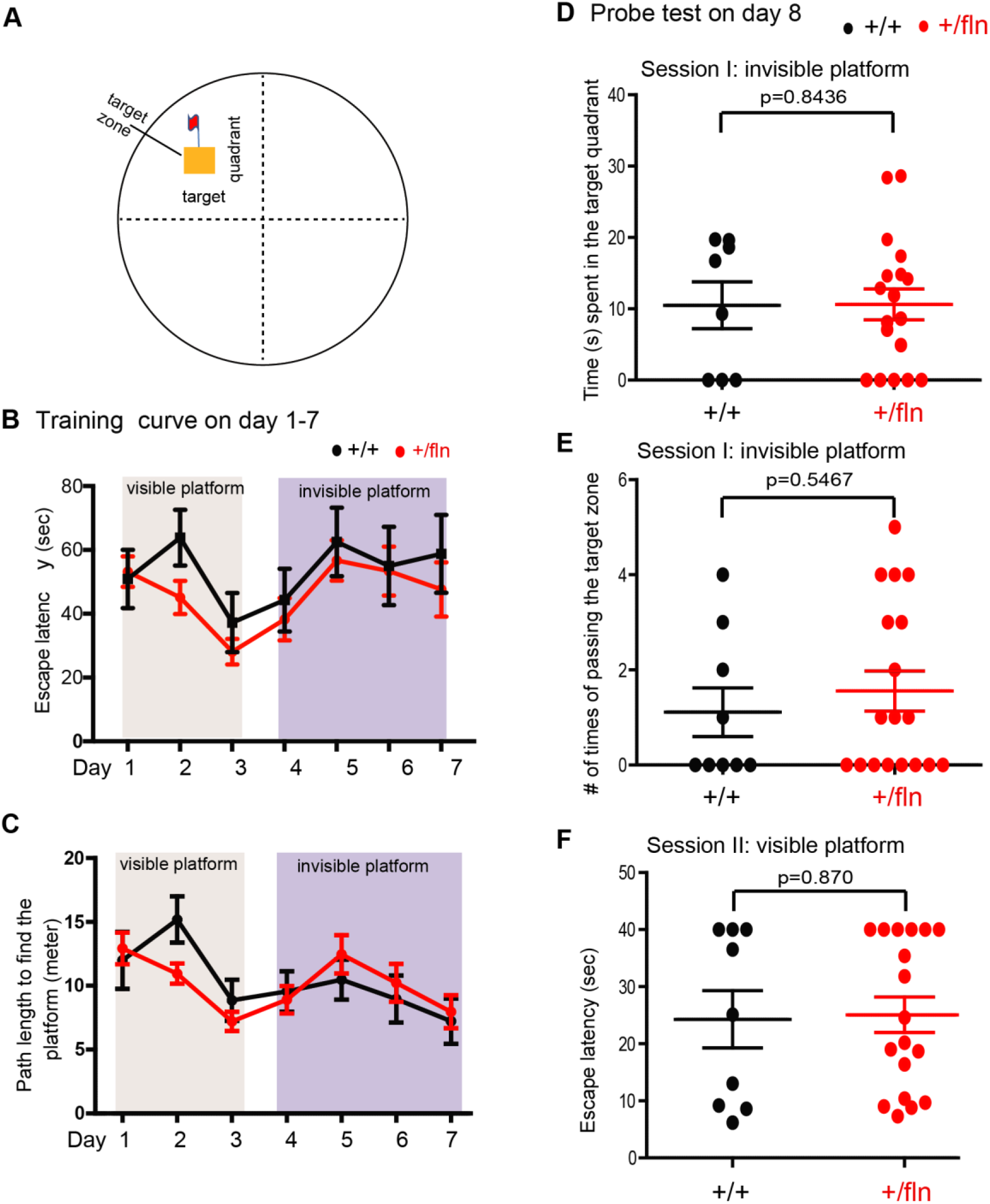
Morris water maze test. (A) Diagram of Morris water maze. The test mice (18 month old) were first trained to find a platform with a visible flag on days 1 to 3 and then a submerged hidden platform on days 4 to 7. The time (B) and path length (C) to find the platform was recorded. On day 8 of probe test, the platform was removed in session I. Times spent by mice in the target quadrant (D) and # passing of the target zone (E) were recorded. In session II, the visible platform was placed back and the latency time (F) was recorded. Mean±SEM, by unpaired t test. NGF^+/+^, n=9, NGF^+/fln^, n=18.

### NGF^R100W^ mutation results in reduction in NGF secretion

To investigate if reduced secretion in NGF could account for the degeneration of peripheral sensory structures and functions, we explored the possibility that R100W mutation impaired the secretion of mature NGF as suggested previously(Carvalho et al., 2011). We dissected and cultured primary fibroblasts of from E18 embryos. Media were collected and the levels of secreted NGF were measured using ELISA. Our data revealed that fibroblasts from either NGF^fln/fln^ (88.32±6.11pg/ml) or NGF^+/fln^ (92.49±11.60 pg/ml) mice secreted significantly less amount of NGF compared to fibroblasts from NGF^+/+^ mice (172±22.06 pg/ml) (Fig 12). This is consistent with our previously published literature showing a significantly lower yield of NGF^R100W^ recombinant protein from 293FT culturing media(Sung et al., 2018). Therefore, reduced secretion of NGF contributes, at least partially, to the peripheral sensory phenotypes as observed in our mouse model.

**Fig 12.**
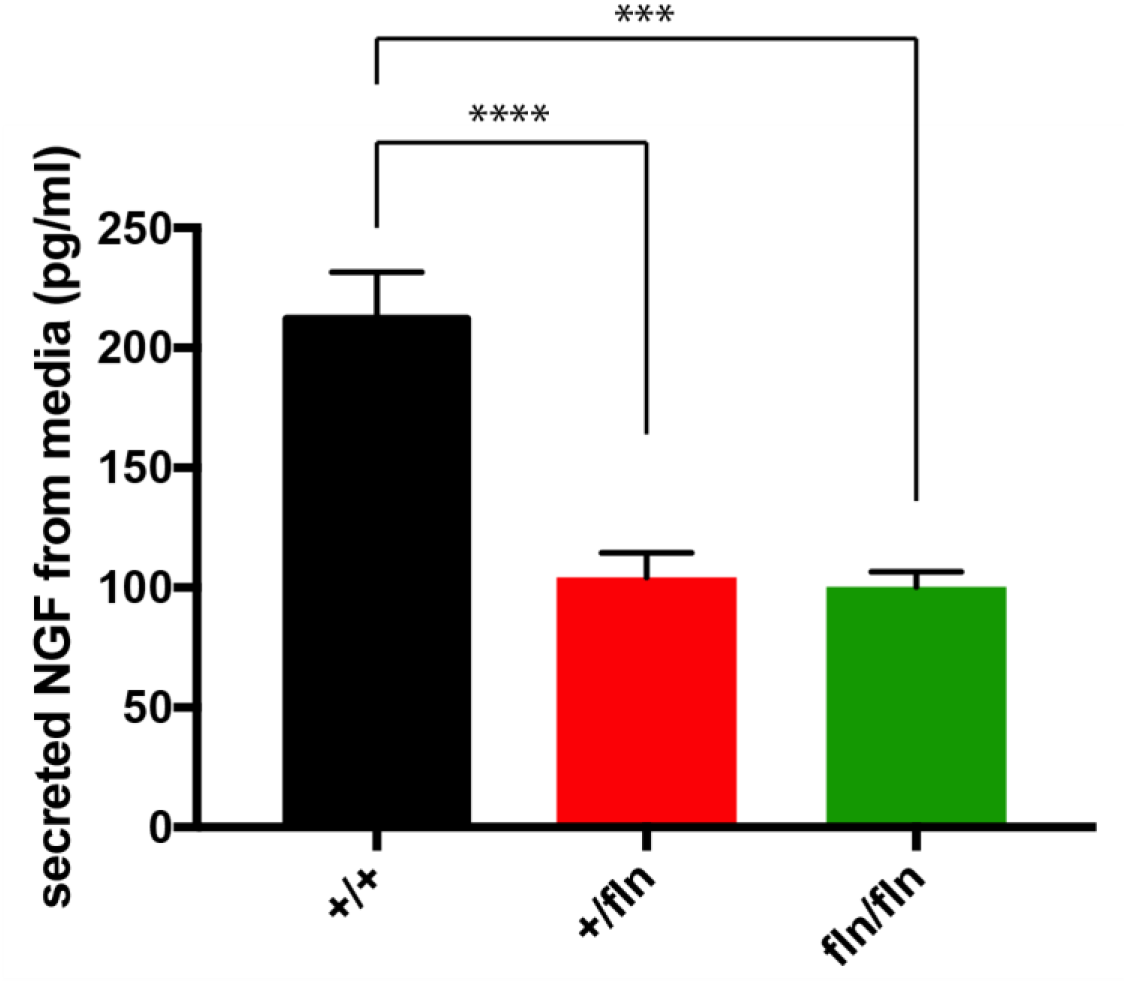
Measurement of NGF secretion in cultures of mouse primary fibroblasts. NGF protein content in media was measured by ELISA. Data from NGF ELISA was presented as mean ± SEM; n=6 for each experimental group.

## Discussion

NGF plays a critical role in supporting the survival and differentiation of specific neuronal populations(Chao, 2003; Huang and Reichardt, 2001; Levi-Montalcini, 1987, 2004; Levi-Montalcini et al., 1995; Levi-Montalcini and Hamburger, 1951; Mobley et al., 1986). Therefore, since its discovery, NGF has been extensively explored for its therapeutic potential for treating neurodegenerative diseases (Apfel, 2002; Apfel et al., 1998; Eriksdotter Jonhagen et al., 1998; McArthur et al., 2000; Olson, 1993; Quasthoff and Hartung, 2001). However, NGF has also been found to induce severe nociceptive responses(Lewin et al., 1993). It is this severe side effect of pain-inducing function that has largely prevented clinical application of NGF.

The naturally occurring NGF^R100W^ mutation in HSAN V patients (Minde et al., 2004b) offers important insight into two important functions of NGF: trophic and nociceptive actions. HSAN V is caused by a missense mutation in NGF gene (661C>T), which leads to a substitution of tryptophan (W) for arginine (R) at position 100 in the mature NGF (NGF^R100W^)(Einarsdottir et al., 2004). Previous clinical study of HSAN V patients found that patients with homozygous mutation had earlier onset and more severe clinical manifestations than heterozygous carriers(de Andrade et al., 2008; Einarsdottir et al., 2004; Minde et al., 2004b). Homozygous patients also reported an inability to detect painful stimuli since birth, which prevents them from feeling pain from heat, bone fractures and joints and leads to destroyed joints in childhood. Sural nerve biopsy with morphometric analysis from homozygous patients further demonstrated a moderate loss of myelinated Aδ fibers and a severer reduction of unmyelinated C fibers(Minde et al., 2004b). These HSAN V patients suffer from severe loss of pain perception but retain normal cognitive function(Capsoni, 2014; Einarsdottir et al., 2004; Minde et al., 2009), suggesting that the NGF^R100W^ mutation may result in selective loss of pain-inducing function, while retaining its trophic function(Capsoni, 2014; Capsoni et al., 2011; Malerba et al., 2015).

To better define the mechanisms of NGF^R100W^ and to study the functional impact of NGF^R100W^ on development, we have generated the first mouse model using the knockin technology. Our study is the first attempt to directly investigate a mice model for NGF^R100W^. We found that the NGF^R100W^ knockin mouse model recapitulates the clinical manifestation of HSAN V; the homozygotes showed severe sensory loss both structurally and functionally at as early as P0, while the heterozygotes developed progressive peripheral sensory neuropathy. Importantly, our tests have revealed that the heterozygotes did not show significant impairment of CNS function. Therefore, our mouse model will serve as an important tool in not only understanding the pathogenesis of HSAN V, but also will be instrumental in uncoupling the nociceptive function from trophic function of NGF.

The NGF^R100W^ homozygotes had significant fewer numbers of small CGRP-positive neurons in their DRGs as well as PGP9.5-IENFs in their skin at birth. They frequently failed to survive to adulthood. The ones that did survived to 6-8 weeks displayed significant deficits in sensing thermal stimuli, a result that appears to be consistent with the lack of sensory fibers and neurons in these mice. In many ways, our NGF^R100W^ homozygotes resemble both the TrkA knockout (Smeyne et al., 1994) and the NGF knockout (NGF^-/-^) mice in their severe sensory phenotypic presentations(Crowley et al., 1994); The NGF knockout mice also failed to respond to noxious mechanical stimuli with profound cell loss of small neurons in their DRGs (Crowley et al., 1994). The similarities between NGF^R100W^ homozygotes and NGF^-/-^ may suggest that the structural and functional deficits in the NGF^R100W^ homozygotes are likely resulted from NGF deficiency. Additional experiments will be needed to confirm if this is indeed the case.

Unlike the homozygotes with early onset of peripheral deficits, NGF^R100W^ heterozygotes did not appear to suffer from any developmental deficits at birth and at 2 months of age. Starting at 9 months, these mice began to show structural degeneration of small CGRP-positive neurons in their DRGs as well as IENFs in their skins. Not surprisingly, they began to develop peripheral sensory deficits. At 18 months of age, these mice showed worsening deficits both structurally and functionally. Therefore, similar to HSAN V patients with a late onset disease, NGF^R100W^ homozygotes show a progressive degeneration of the peripheral sensory system. Intriguingly, the impact of NGF^R100W^ seems to be restricted to the peripheral system, as our examination of the CNS did not identify significant changes both functionally and structurally in these mice.

In some aspects, NGF^R100W^ heterozygotes exhibit phenotypes similar to the NGF hemizygotes (NGF^+/-^), in that these NGF^+/-^ mice also showed a moderate reduction of sensory fiber in skin and CGRP-immunoreactive sensory neurons in the lumbar DRGs(Crowley et al., 1994). Furthermore, these NGF^+/-^ mice displayed decreased responsiveness to pain(Crowley et al., 1994). However, some significant differences exist between the NGF^R100W^ heterozygous mice and the NGF+/-mice; The NGF^+/-^ mice had significantly fewer ChAT-immunoreactive neurons within the medial septal and AChE-positive fibers in CA1, CA3 and dentate gyrus regions of the hippocampal formation(Chen et al., 1997). In addition, NGF^+/-^ mice showed significant acquisition and retention deficits in the water maze test. Whereas in the present study, we did not detect any significant neuronal loss in the hippocampus and media septum of NGF^R100W^ heterozygotes even at the advance age of 18-month old. Furthermore, in a battery of behavioral tests for CNS function ranging from spontaneous alternation, spatial memory, novel object recognition, sociability interaction test and anxiety tests (marble test and light-dark shuttle box test), we did not detect any differences between the NGF^R100W^ heterozygous mice and age-matched +/+ littermate controls. These results demonstrate that NGF^R100W^ show no apparent CNS dysfunction, an important distinction from the TrkA^-/-^ (Smeyne et al., 1994) and the NGF^+/-^ mice(Chen et al., 1997). Our data suggest that the NGF^R100W^ allele in heterozygotes may retain a certain level of trophic function to support the development and function of the brain, while this level of trophic support of the NGF^R100W^ allele is not sufficient in maitaining the structure and function of the peripheral sensory neurons. Taken together, our findings have demonstrated that the NGF^R100W^ heterozygous mice have adult onset, as in HSAN V heterozygous patients, and develop progressive nociceptive deficits and peripheral neuropathy.

The late onset and phenotypic presentations observed in the NGF^R100W^ heterozygous mice are largely in agreement with human HSAN V conditions. The HSAN V heterozygous patients usually had adult onset and a milder progressive course(Einarsdottir et al., 2004; Minde et al., 2004b). For example, one 75-year-old heterozygous carriers underwent a painless fracture of his right ankle at age 48 and 8 years later he developed advanced osteoarthritis in both talocrural joints and left knee with only slight pain(Minde et al., 2004b). The sural nerve biopsies also showed moderate loss of thin myelinated fibers and a severer reduction of unmyelinated fibers. Therefore, the NGF^R100W^ heterozygous mice are an ideal model for studying HSAN V disease.

Recently, human genetic study has linked another case of HSAN V in a consanguineous Arab family to a homozygous mutation in the NGF gene, c.[680C>A]+[681_682delGG] (Carvalho et al., 2011). The mutations reside in the “CGG” codon at the 680-682 position: ‘C” was changed to ‘A’ with “GG” being deleted, resulting in the terminal 15 amino acids of NGF being replaced with a novel 43 amino acid terminal sequence (NGF^V232fs^)(Carvalho et al., 2011). Clinical manifestations of these patients are similar to HSAN IV associated with mutations in TrkA (Indo, 2002), in that they were completely unable to perceive pain, did not sweat, could not discriminate temperature, and had a chronic immunodeficiency (Capsoni, 2014; Carvalho et al., 2011; Indo, 2002). In addition, like HSAN IV patients, these Arabian patients suffered from intellectually disabled(Capsoni, 2014; Carvalho et al., 2011). Therefore, in the case of NGF^V232fs^, it is possible that the mutation results in reduced binding and activation of NGF to TrkA; Alternatively, the mutation impairs the processing of proNGF^V232fs^ and reduces its secretion as demonstrated in in vitro experiments(Carvalho et al., 2011). In either case, TrkA-mediated signaling is expected to be diminished. As a consequence, the development and function of both PNS and CNS are affected.

That the impact of the NGF^R100W^ mutation is largely restricted to PNS function points to some clear distinctions from the NGF^V232fs^ mutation. It is also possible that the NGF^R100W^ mutation has resulted in changes in the processing, secretion and stability of NGF, albeit to a much lesser degree than for NGF^V232fs^. The NGF^R100W^ mutation is located in a highly conserved region that is important for binding and activating p75^NTR^(Einarsdottir et al., 2004). Surface Plasmon Resonance analyses demonstrated that NGF^R100W^ mutation selectively abolishes NGF binding to p75^NTR^, while the affinity for TrkA receptor is retained (Capsoni et al., 2011; Covaceuszach et al., 2010). Therefore, it is likely that the loss of p75^NTR^-mediated signaling in NGF^R100W^ may contribute to impairment of peripheral function in sensing pain. Indeed, there is increasing evidence to support the notion that p75^NTR^ involved in NGF-induced hyperalgesia. Intrathecal administration of antisense oligodeoxynucleotides specific for p75^NTR^ into neuropathic pain model, which reduced its expression in the DRG, attenuated thermal hyperalgesia and mechanical allodynia after nerve injury(Obata et al., 2004). Besides, application of p75^NTR^ blocking antibody inhibited the sensitizing action of NGF to increase the number of action potentials (Zhang and Nicol, 2004) and suppressed mechanical allodynia from crushed sciatic nerve(Fukui et al., 2010). Moreover, administration of neutralizing antibody to p75^NTR^ blocked hyperalgesia arising from an intraplantar injection of NGF(Watanabe et al., 2008). Furthermore, the effect of p75^NTR^ knockout animal mice seemed to be restricted to the sensory nervous system(Lee et al., 1992). This indicates that p75^NTR^ mainly mediate signaling to induce hyperalgesia.

In summary, we have generated a NGF^R100W^ knockin mouse model for studying HSAN V. Our studies have demonstrated that the NGF^R100W^ knockin mice suffer structural and functional degermation of the peripheral sensory system as in HSAN V patients. Furthermore, also as in HSAN V patients, the NGF mutant mice show no significant cognitive deficits, an important feature that distinguishes NGF^R100W^-HSAN V from both NGF^V232fs^-HSAN V and HSAN IV associated with TrkA mutations. This novel mouse model will provide exciting opportunities to further define the disease mechanisms for HSAN V. More importantly, the unique NGF^R100W^ mouse model will facilitate studies to define how NGF^R100W^ retains its trophic signal while the nociceptive function is lost in NGF^R100W^.

## Acknowledgements

We would like to thank Dr William C Mobley for advice and Ms Pauline Hu for her technical assistance; We would also like Ms. Angela Zeng, Mr. Simon Kim for helping with quantitation of IENFs; This research was supported by grants from Tau Consortium (CW); NIH grants AG051848, BX003040, AG051839, AG018440 (RAR).

## Author Contributions

WY, KS, JD and CW designed the study; WY. KS, WX, MJR, ACW, SAS, SF, RKU, SXD, BCG, XO, JR performed the experiments. WY, KS, KJ, NC, RR, JD and CW wrote the paper. All authors discussed the results and commented on the manuscript.

## Supplemental Information

Movie S1. A Movie of PGP9.5-positive IENFs in P0 +/+ mice.

Movie S2. A Movie of PGP9.5-positive IENFs in P0 +/fln mice.

Movie S3. A Movie of PGP9.5-positive IENFs in P0 fln /fln mice.

Movie S4. A Movie of PGP9.5-positive IENFs in 18-month-old +/+ mice.

Movie S5. A Movie of PGP9.5-positive IENFs in 18-month-old +/fln mice.

